# The transcriptomics of phenotypic nonspecificity in *Drosophila melanogaster*

**DOI:** 10.1101/2025.05.13.653887

**Authors:** Gabriella Sidhu, Anthony Percival-Smith

## Abstract

Transcription factor (TF) function is redundant: TF phenotypes are frequently rescued by TFs not resident to the *TF* locus, a phenomenon termed phenotypic nonspecificity. Phenotypic nonspecificity in *Drosophila melanogaster* is not dependent on the DNA binding specificity of the TFs and generally due to genetic complementation. Two TF phenotypes (doublesex [dsx] and apterous [ap]) are rescued by multiple TFs. The rescue by resident TFs, and the rescue and non-rescue by non-resident TFs, of these two phenotypes were used to distinguish between three possible outcomes of the comparison of the TF-dependent mRNA accumulation in these two systems. First, the sets of TF-dependent mRNAs are independent and non-overlapping; second, the sets of TF-dependent mRNAs are independent and overlapping; and third, the sets of TF-dependent mRNAs are constrained and have extensive overlap. The transcriptomes associated with rescue by resident TFs, and rescue and non-rescue by non-resident TFs, of the two TF phenotypes (dsx and ap) provided many examples of extensive overlap indicating regulation of constrained sets of genes. However, the strength of correlation of transcript accumulation observed between the resident and non-resident TFs was not a strong predictor for rescue of the phenotype by the non-resident TFs. The regulation of a constrained set of genes supports the hypothetical assembly of TFs into wolfpacks, and not the hypothetical limited specificity of TF function, as a potential explanation of phenotypic nonspecificity.

**Article Summary:** Transcription factors (TF) control the rate that the information stored in DNA is converted to RNA. Although often considered functionally very specific molecules, TF function is redundant, as frequently the process regulated by one TF is also be regulated by another unrelated TF; a genetic phenomenon termed phenotypic nonspecificity. This paper addresses what patterns of gene expression are brought about by TFs that can regulate, or not regulate, the processes of female abdomen pigmentation and initiation of wing formation. Surprisingly, the patterns of gene expression are very similar. Strangely though, the strength of similarity, or correlation, is not a strong predictor that the TF regulates the process.

## Introduction

The redundancy of transcription factor (TF) function results in the phenomenon of phenotypic nonspecificity where a phenotype due to the lack of expression of the resident TFa expressed from the *TFa* locus is rescued by expression of a non-resident TFb in place of TFa (Percival-Smith 2017; Percival-Smith et al., 2023). In *Drosophila melanogaster*, the rescuing resident and non-resident TFs often do not share similar DNA binding sites nor belong to similar TF families. In most cases of phenotypic rescue, the rescue by the non-resident TFb is due to functional complementation rather than TFb being a downstream/epistatic component of the pathway. In addition, the rescue of the TF phenotypes by the non-resident TFs is differentially pleiotropic.

Phenotypic nonspecificity is a general phenomenon. It is detected at high frequencies with phenotypes induced by the ectopic expression of TFs in Drosophila such as wingless, eyeless, sex-comb-less, ectopic T1 beard formation, female abdominal pigmentation and arista to tarsus transformation phenotypes (Percival-Smith et al., 2005; Percival-Smith 2017; Percival-Smith et al., 2023), and in vertebrates for the re-establishment of pluripotency (Takahashi, and Yamanaka 2006; Maekawa et al., 2011). More importantly, phenotypic non-specificity is detected in 8 of 9 Drosophila *TF* loci tested for rescue (Grieg and Akam 1995; Hirth et al., 1998; Percival-Smith, 2017; Percival-Smith et al., 2023). The estimated frequency of phenotypic nonspecificity of rescue of a TF phenotype in Drosophila is 1 in 10 to 20 TFs rescue. Phenotypic nonspecificity and phenotypic convergence share the conclusion that distinct sets of TFs expressed lead to the same phenotype (Konstantinides et al., 2018). Large scale screens of TFs for the rescue of doublesex (dsx) female abdominal pigmentation pattern and the rescue of the apterous (ap) wingless phenotype identified multiple non-resident TFs that rescue these two phenotypes (Percival-Smith et al., 2023). Here the transcriptomes associated with the expression of resident and non-resident TFs that rescue and do not rescue these two phenotypes are characterized.

The abdominal pigmentation pattern of Drosophila is a major external sexually dimorphic characteristic. The *dsx* gene encodes two functionally distinct TFs that determine secondary sexual characteristics: DSX^M^ determines male characteristics, and DSX^F^ determines female characteristics (Burtis and Baker, 1989). Male abdominal pigmentation does not require DSX^M^ as a *dsx* null mutant has male-like pigmentation. For female abdominal pigmentation, DSX^F^ activates the expression of two functionally redundant TFs, Bric á Brac 1 (BAB1) and BAB2, which suppress the expression of Dopachrome-conversion enzyme encoded by the *yellow* (*y*) gene, and the Beta alanyl dopamine hydrolase encoded by the *tan (t)* gene (Couderc et al., 2002; Han et al., 2002; True et al., 2005; Williams et al., 2008; Camino et al., 2015; Roeske et al., 2018). The male-like abdominal pigmentation of the dsx^null^ phenotype is rescued to female-like pigmentation by the expression of DSX^F^, BAB1, Antennapedia (ANTP), Buttonhead (BTD), Eyelesss (EY), Hunchback (HB), Knirps (KNI), Scalloped (SD), Sine ocullus (SO) or Oddskipped (ODD) but not by the expression of Forked head box subgroup O (FOXO) or Squeeze (SQZ) (Percival-Smith et al., 2023). The rescue by expression of BAB1 is expected as it functions downstream of DSX^F^; however, the other TFs that rescue have no normal requirement in abdominal pigmentation pattern. Analysis of the epistatic interactions between expression of DSX^F^, BAB1, ANTP, EY or ODD and *dsx^null^*or *bab* alleles suggests that BAB1, ANTP, EY and ODD all function after DSX^F^. As BAB1 and 2 bind *cis*-regulatory elements in *y* and likely *t* repressing expression, BAB1, ANTP, EY and ODD likely function as interchangeable repressors of these two genes (Roeske et al., 2018). DSX^F^ activates BAB1 and BAB2 expression at around 72 hours after pupa formation (APF), and the BAB dependent repression of a *y::GFP* transcriptional fusion gene is detected at 85-95 h APF, the P13-15 pupal stages (Williams et al., 2008; Roeske et al., 2018).

The spatially restricted expression of the Apterous (AP) TF during the second instar larval stage establishes the dorsal compartment of the wing imaginal disc, and initiates a pathway required for wing formation (Cohen et al., 1992; Fisher et al., 2017). At the late first/early second instar larval stage, the first step in determination of the dorsal ventral axis of the wing imaginal disc is mutual negative interactions between Vein (VN) expressed on the future dorsal side and Wingless (WG) expressed on the future ventral side of the wing imaginal disc (Couso et al., 1993; Simcox et al., 1996). During the second instar larval stage, the VN signal activates the dorsal expression of AP that establishes the dorsal compartment. For wing formation, AP activates the expression of Serrate (SER) and Fringe (FRN) (Diaz-Benjumea and Cohen 1993; Diaz-Benjumea and Cohen 1995; Irvine and Wieschaus, 1994). SER and FRN regulate the activity of the Notch pathway that then activates WG expression in the cells of the future wing margin (Klein and Arias, 1998). WG expression along the wing margin is established by the late second/early third instar larval stage (Ng et al., 1996). Subsequently WG and Decapentaplegic (DPP) promote the growth of the wing pouch, which during the pupal stages differentiates into the adult wing (Zecca and Struhl, 2021). The wingless phenotype of an *ap^null^*mutant is rescued by the expression of AP, Caudal (CAD), Tramtrack (TTK) or Myb oncogene-like (MYB) but not Sisterless A (SISA). CAD and TTK are not required for wing development but loss of MYB does result in an altered wing suggesting a possible requirement in wing formation (Percival-Smith et al., 2023).

This paper characterizes the transcriptomes associated with the expression of the TFs BAB1, ANTP, EY, FOXO, ODD, or SQZ in a *dsx^null^* mutant background and the transcriptomes associated with the expression of the TFs AP, CAD, MYB, SISA or TTK in an *ap^null^* mutant background. In these two systems (dsx and ap), the TF-dependent mRNA accumulations associated with the expression of various TFs were compared to distinguish between three potential outcomes. First, the rescuing resident and non-resident TFs regulate completely independent, non-overlapping sets of genes; second, the TFs regulate independent overlapping sets of genes; or third, the TFs co-regulate a constrained set of genes. The TFs regulate expression of non-independent, overlapping sets of mRNAs supporting the third outcome: the regulation of a constrained set of genes. These results support the proposal of the assembly of TFs into membrane-less compartments (wolfpacks) as a potential explanation of phenotypic nonspecificity (Staller 2022; Percival-Smith et al., 2023).

## Materials and methods

### Drosophila husbandry

Flies were maintained at 24 C and 60% humidity. The flies were reared in 20ml vials containing corn meal media [1% (w/v) Drosophila grade agar, 6% (w/v) sucrose, 10% (w/v) cornmeal, 1.5% (w/v) yeast, and 0.375% (w/v) 2-methyl hydroxybenzoate]. For collection of early third instar larvae, flies laid eggs on apple juice plates [2.5% (w/v) Drosophila grade agar, 6% (w/v) sucrose, 50% apple juice and 0.3% (w/v) 2-methyl hydroxybenzoate] smeared with a patch of yeast paste on the first day, and the progeny were allowed one day to hatch. After hatching, a layer of mashed corn meal media mixed with yeast paste was spread across the plate. On the fourth day, early third instar larvae (0-8 hours old) were present for collection. For collection of P13-15 pupa, vials with wandering third instar larvae were inspected for white prepupa whose positions were marked on the vial with a permanent marker. The vials were incubated for four days and P13-15 pupa were identified based on described markers (Ashburner, 1989).

### The Drosophila crosses and genotype identification

The stocks used for the experiments in the *dsxGAL4/dsx^1^*null background were generated previously (Robinett et al., 2010; Percival-Smith et al., 2023; TableS1). The following crosses were performed: *y w; dsx^1^/TM6B, Tb, P{walLy}* X *w; dsxGAL4/TM6B, Tb,* and *y w; P{UASX, w^+^}; dsx^1^/TM6B, Tb, P{walLy}* (where *X* could be *Antp, bab1, dsx^M^, ey, foxo*, or *sqz*) X *w; dsxGAL4/TM6B, Tb*, and *y w; P{UASdsx^F^, w^+^}, dsx^1^/TM6B, Tb, P{walLy}* X *w; dsxGAL4/TM6B, Tb*. The non-Tb pupae in the progeny were *dsxGAL4/dsx^1^*and expressing GAL4 alone or expressing GAL4 with a TF.

To identify wild-type male and female pupae, wandering third instar larvae were separated based on the size of the larval gonads: male gonads are large causing a characteristic clearing (hole) of the opaque fat bodies. The male and female larvae were allowed to develop for four days in separate vials.

The stocks used for the experiments in the *apGAL4/ap^MIO1996-FLPSTOP.D^* null background were assembled using standard Drosophila crossing schemes (Table S1). The following crosses were performed: *y w; apGAL4/CyO, P{UbiGFP, w^+^}PAD1* X *y w; ap^MIO1996-FLPSTOP.D^ / CyO, P{UbiGFP, w^+^}PAD1*, and *y w; apGAL4/CyO, P{UbiGFP, w^+^}PAD1 X y w; ap^MIO1996-FLPSTOP.D^ / CyO, P{UbiGFP, w^+^}PAD1; P{UASap, w^+^}*, and *y w; apGAL4/CyO, P{UbiGFP, w^+^}PAD1 X y w; ap^MIO1996-FLPSTOP.D^ / CyO, P{UbiGFP, w^+^}PAD1; M{UASX, GW}ZH-86Fb* (where *X* is *cad, ttk* or *sisA* fused to *3XHA tag*, or *myb*). The non-GFP-expressing larvae in the progeny were *apGAL4/ap^MIO1996-FLPSTOP.D^* expressing GAL4 alone or expressing GAL4 along with a TF.

### Tissue dissection

For dissection of P13-15 abdomens, pupae of the correct genotype were identified and the dissected from the pupal case. In Drosophila Ringer’s solution (3mM CaCl_2_, 182mM KCl, 46mM NaCl, 10mM Tris HCl pH 7.2), the abdomen was dissected by grabbing the pupa between the thorax and abdomen with a pair of forceps and cutting off the abdomen by sliding a second pair of forceps along the first. The abdomens were placed in a microcentrifuge tube containing Ringer’s solution on ice.

For dissection of late third instar larval wing imaginal discs, wandering third instar larvae were placed in Ringer’s solution and screened for larvae that lacked GFP expression (no *CyO* balancer). The larvae were dissected using sharp forceps by grabbing the larvae in the middle with the first pair of forceps and pulling up a strip of dorsal cuticle with the second pair. The wing imaginal discs were identified and picked up with forceps and stored in a microcentrifuge tube containing Ringer’s solution on ice.

Early third instar larvae (0-8 hours after molting) were differentiated from late second instar larvae by the distinct, stage specific, structure of the anterior spiracles (Ashburner, 1989). For dissection of early third instar larval tissue, larvae were placed in Ringer’s solution and screened for lack of GFP expression. The larvae were dissected using sharp forceps by grabbing the larvae in the middle and pulling up a strip of dorsal cuticle to the mouth hooks and then cutting the larvae in half by sliding one pair of forceps over the other. The head skeleton including the attached gut and brain were removed with forceps leaving a patch of anterior cuticle that had the early third instar leg, wing and haltere imaginal discs attached. The dissected patches were stored in a microcentrifuge tube containing Ringer’s solution on ice.

### RNA extraction and RNAseq

The P13-15 abdomen samples had 5-8 abdomens per sample; the late third instar larval wing disc samples had 20-35 dissected wing discs per sample; and the early third instar larval anterior samples had 8-12 anterior cuticle patches per sample. Total RNA was extracted from the tissues using the RNeasy mini kit (QIAGEN). The number of biological replicates (≥5) were chosen based on published recommendations (Conesa et al., 2016) and was met for 20 sets of replicates with the exception of n=4 for 4 sets of replicates (see Table S2). The RNA samples were sequenced by Inomomic Inc. (Shenzhen, China) using DNBSEQ Eukaryotic Strand-specific Transcriptome Resequencing PE100 in five separate batches and PE150 for the final sixth batch. The number of clean forward reads and their batch and method are given in Table S2.

### RNA seq data analyses

The reads were aligned to the BDGP6 assembly of the *Drosophila melanogaster* genome using HISAT2 (version 2.2.1) (Kim et al, 2019). The count data was extracted with featureCounts (version 2.0.1) with the BDGP6.46.110 genome assembly (Liao et al., 2014) from the file of forward reads. Differential mRNA accumulation, often referred to as differentially expressed genes (DEG), was detected with DEseq2 (version 1.40.2) (Love et al., 2014) for a sample size of ≥5 (with 4 noted exceptions of n=4), based on the recommendation of Li et al., 2022.

The base mean threshold for defining expressed genes from a DEseq2 analysis was chosen to be 1 rather than 10 because DEseq2 identified DEGs (*Padj*<0.05) between base means of 1 and 10, but not 0 and 1, and for the question posed, analysis of all DEGs needed to be included because the question pertains more to the structure of the TF-dependent transcriptome than the functional importance of individual genes identified. This is also the reason that no effect size threshold filter was applied to the data as well. Because the data analyzed was collected in six batches of sequencing, the principle components of variation in the DEseq2 analyses was determined (Figure S1). Generally for the dsx set of experiments, the PC2 (5-20 percent of the variation) showed separation of the control and TF samples and the PC1 (60-80 percent of the variation) showed the variation between replicates and the individual batches were spread out across PC1 in the dsx set of experiments suggesting a lack of a batch effect in batches 1, 2 and 3. The ANTP PCA was an exception showing a clustering of the TF from the control sloped across both PC1 and 2, and the ANTP analysis had the highest number of DEGs in the dsx set. The ap sets of experiments were unbalanced and had very high numbers of DEGs. For the L3 set of data the TF and control were clearly separated along PC1, but due to the unbalanced origin of the samples the effect of the TF could not be discerned from an effect of batch along PC1. The separation of the TF and control data occurred in 5 of the 6 PCAs of the E3 data along PC1 but to a lesser degree than L3; however, in the CAD analysis at E3, the data for the three batches was dispersed across PC1 (separation of the TF and control data was along PC2) suggesting a lack of a batch effect in batches 4, 5 and 6. Therefore, the data was analyzed in DEseq2 without taking a batch effect into consideration to reduce the degrees of freedom for the analyses. When plotting the log_2_ fold changes of two TF-dependent transcriptomes, the gene data with an NA (filtered out by DEseq2) for one of the two TFs were not included on the graphs.

Venn diagrams were generated with VennDiagram package version 1.7.3 (Chen and Boutros 2011). For the expected mean of the overlap of gene expression between *a* and *b* based on independent regulation gene expression was calculated by multiplying the fractions, DEG*a*/Total expressed genes*a* X DEG*b*/total expressed genes*b*, and multiplying the resulting fraction by the average of total expressed genes of *a* and *b*. The Monte Carlo simulation of the number of overlapping differentially expressed genes between two or more analyses was performed in two steps. First a random number was chosen from a set of unique numbers equal to the total number of expressed genes determined in a DEseq2 analysis (base mean>1) (Supplemental method S1). This was repeated until the total number of unique numbers chosen was equal to the number of DEGs identified in the same DEseq2 analysis. These sets of numbers representing DEGs were generated for two or more independent DEseq2 analyses. In the next step, the union between the sets, which are the numbers shared between the sets, were counted. These Monte Carlo simulations were repeated 1000 times generating a list of 1000 simulated, shared DEG overlaps. The 95% confidence interval for each TF comparison was defined by the 25^th^ numbers from both the lowest and highest numbers of DEGs generated in the 1000 simulations.

For estimating the number of transcripts expressed from the endogenous *TF* loci and *UAS TF* constructs, Salmon (version 1.10.1) was used (Patro et al., 2017). Salmon was provided with the BDGP6 assembly of the *Drosophila melanogaster* transcriptome supplemented with FASTA files of the UAS constructs used (Supplemental data S1). The count data was then normalized to the sample with the highest total counts. In addition, because the relative lengths of transcripts affect the number of counts detected, the count data were divided by the length of the sequence used to search the reads in Salmon (counts/kb).

The virtual in situs were generated with the method described in Everetts et al., 2021. The script used to generate virtual in situs from a list of genes is included in the supplemental method S2. The virtual in situs were assigned to one of 11 expression categories by visual inspection.

### Statistical analysis

When DEseq2 performs an analysis of two sets of samples, the data is normalized; however, when the normalized count data of the same control samples of two DEseq2 analyses are compared, they differ. To avoid this variation in control count data when comparing multiple analyses, the featureCounts data were normalized to the one with the highest total counts in the dataset and the data plotted.

For the statistical analysis, the count data was treated as continuous data because all the counts of interest were high and above zero. The data were log transformed for some analyses to meet the criteria of an ordinary ANOVA, and multiple pairwise comparisons *post hoc* analyses were performed with Dunnett’s test. For some analyses a non-parametric analysis was required. These analyses were performed in the Prism statistics package (Graphpad).

### *Wingless* in situ hybridization

Late third instar larvae were dissected by grabbing the larva in the middle and ripping a strip of dorsal cuticle to the anterior tip and cutting the larvae in half. The tissue was fixed for 30 min in 5% formaldehyde in 1X Phosphate buffed saline (PBS). The fixative was washed off with repeated washes of 1XPBS and the samples stared in 100% methanol. Thirty-five *wg* primers with the structure 21 *wg* bases followed by 30 Gs were synthesised. The 35 primers were mixed together at equal molarity and hybridized with a 3’ Alex488 labeled C15 primer at a molar ratio of 1-3.5, respectively. The tissue was hybridized at a concentration of 100nM overnight at 37 C and repeatedly washed as described in Calvo et al., 2021. The tissue was dispersed with a sharp hypodermic needle, mounted in Vectashield vibrance and the nuclei and *WG* imaged on a Nikon (Eclipse Ti2E) inverted microscope.

## Results

### Expression levels of the *UASTF* constructs

To assess whether the *UASTF* constructs were over- or under-expressed relative to the endogenous *TF* locus, the program Salmon was used to estimate the mRNA counts originating from the *UASTF* transgenes and the endogenous *dsx* and *ap* loci by adding the sequences of the *UASTF* transcripts to the Drosophila transcriptome (Patro et al., 2017). The transgenes, with one exception, were expressed higher relative to the endogenous loci: transcripts from *UASdsx^F^* accumulate 1.8 times higher than endogenous *DSX^F^*transcripts, and transcripts from *UASap* accumulate 3.5 times higher in late third instar larval wing discs than transcripts from the endogenous *ap* locus (Table 1).

**Table 1.**
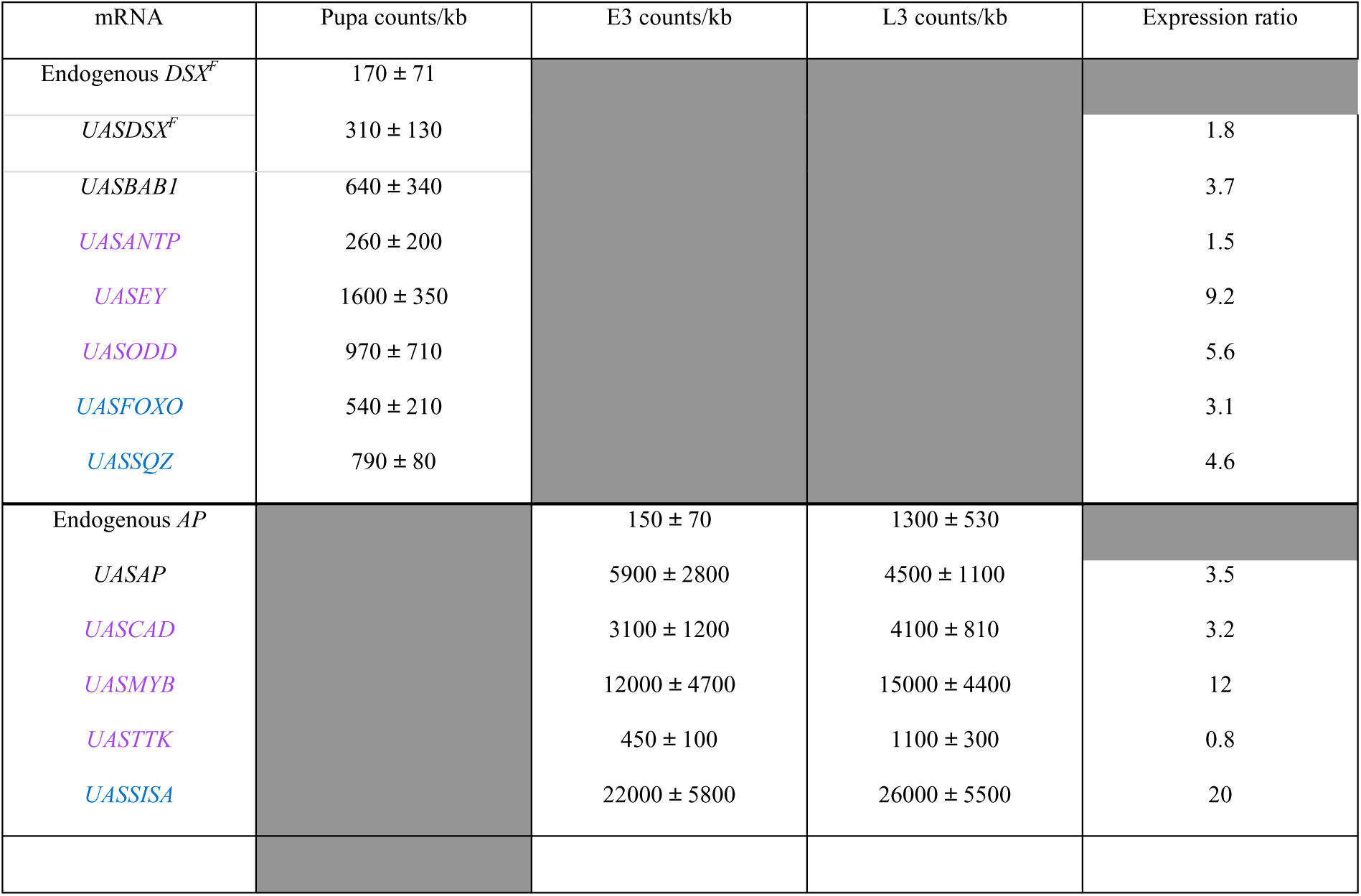
Transcript abundance of expression from the *UASTF* transgenes relative to the endogenous locus.

### DSX^F^ and BAB1 dependent mRNA accumulation

One expectation of DSX^F^ activating BAB expression for female pigmentation is that DSX^F^ expressed in a *dsx^null^* mutant should regulate a large set of genes that includes both DSX^F^ and BAB regulated genes; whereas, when BAB1 is expressed in a *dsx^null^* mutant, only the smaller subset of BAB1 regulated genes should be expressed. In a Venn diagram of the DSX^F^ and BAB1 dependent mRNAs, the set of BAB1 dependent mRNAs is smaller and 57% these mRNAs overlap with the DSX^F^ dependent mRNAs (base mean >1; *P_adj_* <0.05). Plotting the log_2_ fold change (L2FC) of the mRNAs displayed in the Venn diagram when DSX^F^ is expressed on the x-axis and fold change of the mRNAs when BAB1 is expressed on the y-axis showed a strong correlation of the TF dependent mRNAs shared by these TFs (red/burgundy points; R^2^ =0.88, slope=1.07 y-intercept= 0.06). Of the 504 mRNAs dependent on both DSX^F^ and BAB1 in the Venn diagram only 7 showed alternate regulation (higher accumulation than the control with one TF; lower with the other TF). A number of mRNAs (estimated at 80-150 mRNAs) that were not identified in the DSX^F^ set (green) and not identified in the BAB1 set (blue) showed similar fold changes in gene expression with both TFs, and this is due to not meeting the *P_adj_* <0.05 threshold with one of the TFs indicating that the 504 mRNAs dependent on both DSX^F^ and BAB1 is potentially an underrepresentation. 343 DSX^F^-dependent mRNAs (light blue points close to the x-axis) showed a less than two-fold change when BAB1 was expressed, as expected if there are DSX^F^ regulated genes that are not regulated by BAB1. However, there were 168 examples of BAB1-dependent mRNAs that showed a less than two-fold change with DSX^F^ (light green points close to the y-axis). Overall, the expectation of the set of BAB1-dependent mRNAs being nested in the set of DSX^F^-dependent mRNAs is generally supported.

### TF-dependent transcriptomes associated with rescue/non-rescue of female abdominal pigmentation

The TFs DSX^F^, BAB1, ANTP, EY, and ODD rescue female abdominal pigmentation of *dsx^null^* mutants and the TFs FOXO and SQZ do not rescue female pigmentation (Figure 2a; Percival-Smith et al., 2023). From analysis of epistasis, the BAB1, ANTP, EY and ODD TFs act after DSX^F^ and likely complement the loss of BAB expression in the *dsx^null^* mutant. The BAB1, ANTP, EY, ODD, FOXO and SQZ dependent mRNAs expressed in the *dsx^null^* abdomens of P13-15 pupa were identified. Venn diagrams have an overlapping pattern of sharing between the sets of TF-dependent mRNAs with 3 mRNAs shared by BAB1, ANTP, EY and ODD (Figure 2c). A similar overlapping pattern of sharing was observed when SQZ and FOXO were added (Figure 2d). Although this may suggest that the TFs potentially regulate independent sets of genes, the data was analyzed in two ways to assess further whether this pattern is truly stochastic and meets the criterion of independence.

**Figure 1.**
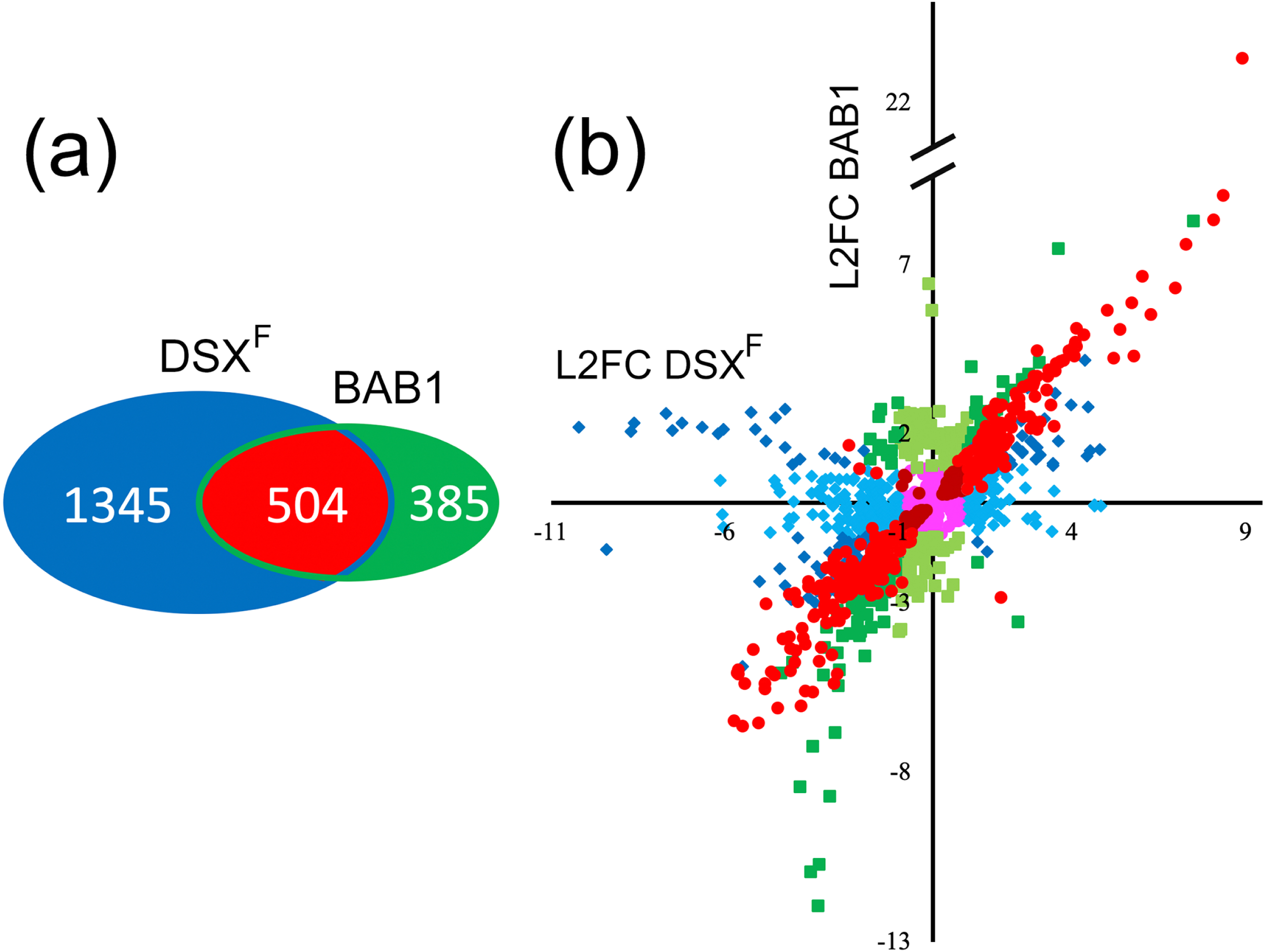
Analyses of the DSX^F^ and BAB1 dependent transcriptomes. a A Venn diagram of the DEGs of the DSX^F^ and BAB1 dependent transcriptomes. b Scatter plot of the log_2_ fold change (L2FC) of DSX^F^-dependent mRNA accumulation (blue and red points) plotted on the x-axis versus BAB1-dependent mRNA accumulation (green and red points) plotted on the y-axis. The circular points in red and burgundy are DEGs identified in both the DSX^F^ and BAB1 transcriptomes (*P_adj_* < 0.05). The red points have a L2FC >1 and <-1; the burgundy points have a L2FC < 1 and > -1. The diamond blue/light blue points are DEGs identified in the DSX^F^ transcriptome (*P_adj_* <0.05) but not the BAB1 transcriptome (*P_adj_* >0.05). The blue points have a L2FC >1 and < -1 with BAB1; light blue have a L2FC <1 and > -1 with BAB1. The square green points are DEGs identified in the BAB1 transcriptome (*P_adj_* < 0.05) and not the DSX^F^ transcriptome (*P_adj_* < 0.05). The green points have a L2FC >1 and <-1 with DSX^F^; light green L2FC < 1; >-1 with DSX^F^. Magenta points were identified as DEGs in either the DSX^F^ or BAB1 transcriptomes with a L2FC of <1 and ώ-1 in both.

**Figure 2.**
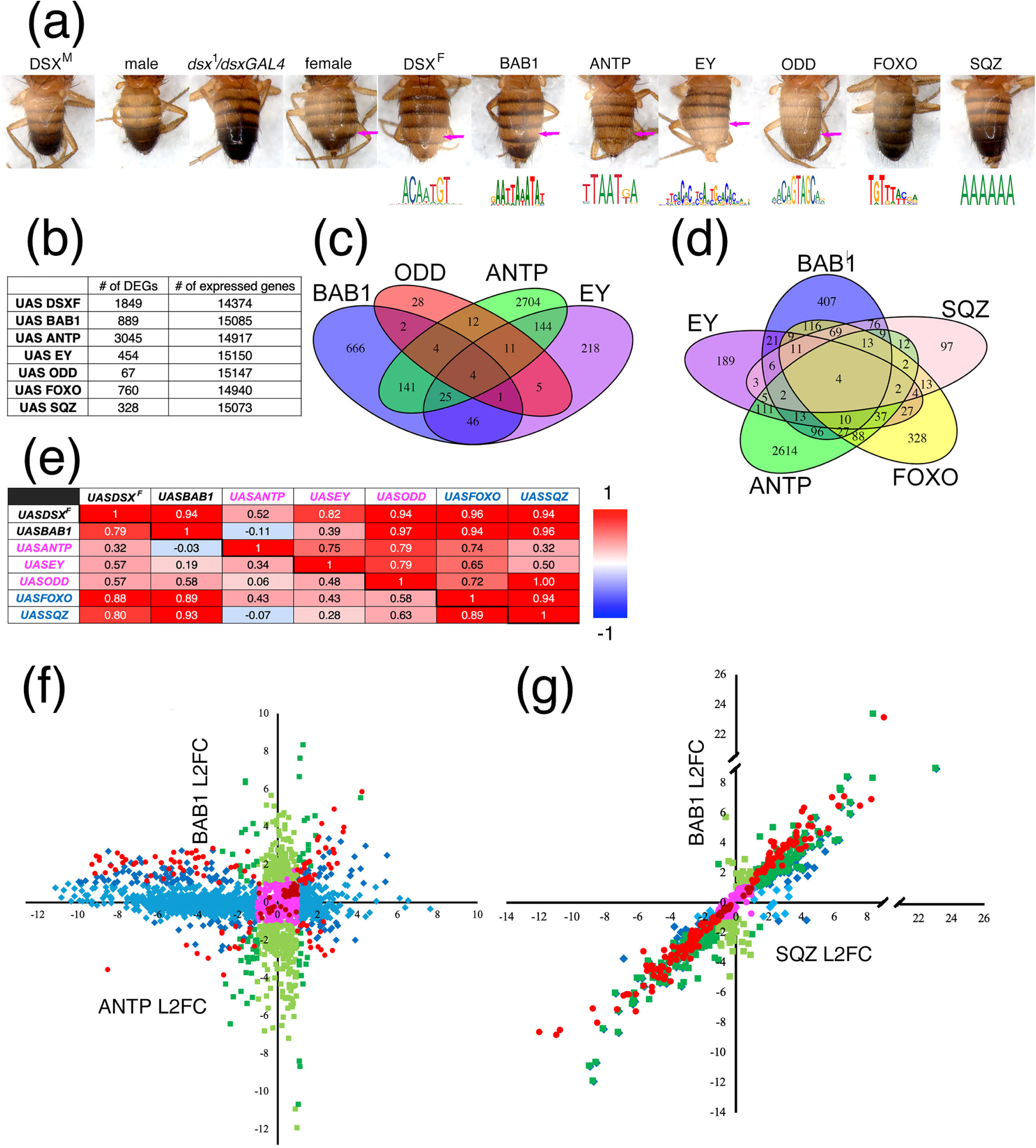
Analysis of the transcriptomes associated and not associated with rescue of female pigmentation in a *dsx^1^/dsxGAL4* null mutant background. a. Rescue of female pigmentation by expression of DSX^F^, BAB1, ANTP, EY or ODD but not by FOXO or SQZ in a *dsx^1^/dsxGAL4* mutant background. Below the image of the abdominal phenotype associated with expression of a TF is a sequence logo of the DNA binding site recognized by the TF (Vliegle et al., 2006). The pink arrows indicate depigmentation of tergite 5. b. Table of the number of DEGs and expressed genes (base mean>1) identified in the DEseq2 analyses. c. Venn diagram of the DEGs of the BAB1, ANTP, EY and ODD transcriptomes. d. Venn diagram of the DEGs of the BAB1, ANTP, EY, FOXO and SQZ transcriptomes. e Table of Pearson’s correlations between DEGs of various TF transcriptomes. The diagonal separates two sets of analyses. The Pearson’s correlations below and left of the diagonal are for all the DEGs identified in two TF-dependent transcriptomes (all the points [red, burgundy, green, blue, magenta] in panels f and g). The Pearson’s correlations above and right of the diagonal are for the DEGs shared between both TF-dependent transcriptomes (the red and burgundy points in panels f and g). f. Scatter plot of the log_2_ fold change (L2FC) of ANTP dependent mRNA accumulation plotted on the x-axis versus BAB1 dependent mRNA accumulation plotted on the y-axis. The circular points in red and burgundy are DEGs identified in both the ANTP and BAB1 transcriptomes (*P_adj_* < 0.05). The red points have a L2FC >1 and <-1; the burgundy points have a L2FC < 1 and > -1. The diamond blue/light blue points are DEGs identified in the ANTP transcriptome (*P_adj_* <0.05) but not the BAB1 transcriptome (*P_adj_* >0.05). The blue points have a L2FC >1 and < -1 with BAB1; light blue have a L2FC <1 and > -1 with BAB1. The square green points are DEGs identified in the BAB1 transcriptome (*P_adj_* < 0.05) and not the ANTP transcriptome (*P_adj_* > 0.05). The green points have a L2FC >1 and <-1 with ANTP; light green L2FC < 1; >-1. Magenta points are DEGs in either the ANTP or the BAB1 transcriptomes with a L2FC of <1 and >-1 in both. g. Scatter plot of the log_2_ fold change (L2FC) of SQZ dependent mRNA accumulation plotted on the x-axis versus BAB1 dependent mRNA accumulation plotted on the y-axis. The points are organized in the same way as panel f with SQZ being substituted for ANTP.

First, the number of DEGs and total expressed mRNAs were used to repeatedly simulate the expected number of genes that would overlap to derive the 95% confidence interval (CI) for the expected values. The observed values were then compared with the simulated data to assess whether they fall within the CI expected for stochastic behaviour (Table 2). With the exception of 2 of 54 comparisons (BAB1 and ANTP; ANTP and SQZ), the observed number was higher than the simulated mean, the 52 observed numbers did not fall in the simulated 95% CI and the difference between the observed number and simulated mean have high z-scores indicating non-independence (z-score = the difference between the observed number and the simulated mean/simulated standard deviation).

**Table 2.** Non-independence of the overlap of transcriptomes in the rescue and non-rescue of dsx.

| TF comparison | Shared genes | Expected | 95% CI<sup>a</sup> | Simulated | z-score |
| --- | --- | --- | --- | --- | --- |
| BAB, ANTP | 174* | 180 | 157 - 202 | 179 | 0 |
| BAB, ODD | 11 | 4 | 1 - 8 | 4 | 4 |
| BAB, EY | 76 | 27 | 17 - 37 | 27 | 10 |
| BAB, SQZ | 190 | 19 | 11 - 28 | 19 | 43 |
| BAB, FOXO | 259 | 45 | 33 - 57 | 45 | 36 |
| ANTP, ODD | 31 | 14 | 7 - 20 | 13 | 6 |
| ANTP, EY | 184 | 92 | 75 - 108 | 91 | 10 |
| ANTP, SQZ | 49 | 67 | 52 - 82 | 66 | -2 |
| ANTP, FOXO | 183 | 155 | 135 - 177 | 155 | 3 |
| ODD, EY | 21 | 2 | 0 - 5 | 2 | 19 |
| ODD, SQZ | 7 | 1 | 0 - 4 | 1 | 6 |
| ODD, FOXO | 22 | 3 | 0 - 7 | 3 | 9 |
| EY, SQZ | 37 | 10 | 4 - 16 | 10 | 9 |
| EY, FOXO | 104 | 23 | 15 - 32 | 23 | 20 |
| SQZ, FOXO | 118 | 17 | 10 - 24 | 17 | 25 |
| BAB, ANTP, ODD | 8 | 1 | 0 - 3 | 0.8 | 7 |
| BAB, ANTP, EY | 29 | 5 | 1 - 11 | 5 | 12 |
| BAB, ANTP, SQZ | 28 | 4 | 1 - 8 | 4 | 12 |
| BAB, ANTP, FOXO | 54 | 9 | 4 - 15 | 9 | 15 |
| BAB, ODD, EY | 5 | 0.1 | 0 - 1 | 0.1 | 12 |
| BAB, ODD, SQZ | 3 | 0.1 | 0 - 1 | 0.1 | 10 |
| BAB, ODD, FOXO | 8 | 0.2 | 0 - 1 | 0.2 | 19 |
| BAB, EY, SQZ | 23 | 1 | 0 - 2 | 0.6 | 28 |
| BAB, EY, FOXO | 34 | 1 | 0 - 4 | 1 | 33 |
| BAB, SQZ, FOXO | 97 | 1 | 0 - 3 | 0.9 | 96 |
| ANTP, ODD, EY | 15 | 0.4 | 0 - 2 | 0.4 | 15 |
| ANTP, ODD, SQZ | 4 | 0.3 | 0 - 2 | 0.3 | 4 |
| ANTP, ODD, FOXO | 13 | 1 | 0 - 3 | 0.7 | 12 |
| ANTP, EY, SQZ | 13 | 2 | 0 - 5 | 2 | 11 |
| ANTP, EY, FOXO | 53 | 5 | 1 - 9 | 5 | 24 |
| ANTP, SQZ, FOXO | 21 | 3 | 0 - 8 | 3 | 9 |
| ODD, EY, SQZ | 4 | 0.04 | 0 - 1 | 0.05 | 20 |
| ODD, EY, FOXO | 13 | 0.1 | 0 - 1 | 0.1 | 43 |
| ODD, SQZ, FOXO | 6 | 0.1 | 0 - 1 | 0.1 | 20 |
| EY, SQZ, FOXO | 21 | 0.5 | 0 - 2 | 0.5 | 29 |
| BAB, ODD, EY, ANTP | 4 | 0.02 | 0 - 1 | 0.03 | 20 |
| BAB, ODD, EY, FOXO | 4 | 0.01 | 0 - 0 | 0.01 | 40 |
| BAB, ODD, EY, SQZ | 2 | 0.003 | 0 - 0 | 0.002 | 50 |
| BAB, ODD, ANTP, FOXO | 6 | 0.04 | 0 - 1 | 0.04 | 30 |
| BAB, ODD, ANTP, SQZ | 3 | 0.02 | 0 - 0 | 0.02 | 30 |
| BAB, ODD, FOXO, SQZ | 3 | 0.004 | 0 - 0 | 0.01 | 30 |
| BAB, EY, ANTP, FOXO | 14 | 0.3 | 0 - 2 | 0.3 | 27 |
| BAB, EY, ANTP, SQZ | 6 | 0.1 | 0 - 1 | 0.1 | 20 |
| BAB, EY, FOXO, SQZ | 15 | 0.03 | 0 - 1 | 0.03 | 75 |
| BAB, ANTP, FOXO, SQZ | 17 | 9 | 0 - 1 | 0.2 | 34 |
| ODD, EY, ANTP, FOXO | 10 | 0.02 | 0 - 0 | 0.02 | 100 |
| ODD, EY, ANTP, SQZ | 3 | 0.01 | 0 - 0 | 0.02 | 30 |
| ODD, EY, FOXO, SQZ | 4 | 1 | 0 - 0 | 0.002 | 100 |
| ODD, ANTP, FOXO, SQZ | 4 | 0.02 | 0 - 0 | 0.01 | 40 |
| EY, ANTP, FOXO, SQZ | 6 | 0.1 | 0 - 1 | 0.1 | 20 |
| BAB, ODD, EY, ANTP, FOXO | 3 | 0.001 | 0 - 0 | 0.001 | 100 |
| BAB, ODD, EY, ANTP, SQZ | 2 | 0.001 | 0 - 0 | 0.002 | 50 |
| BAB, ODD, EY, FOXO, SQZ | 2 | 0.0001 | 0 - 0 | 0.002 | 50 |
| BAB, ODD, ANTP, FOXO, SQZ | 3 | 0.001 | 0 - 0 | 0 | 3388 |
| BAB, EY, ANTP, FOXO, SQZ | 4 | 0.01 | 0 - 0 | 0.003 | 40 |
| ODD, EY, ANTP, FOXO, SQZ | 3 | 0.0005 | 0 - 0 | 0 | 6657 |
| BAB, ODD, EY, ANTP, FOXO, SQZ | 2 | 0.00003 | 0 - 0 | 0 | 75274 |

^ CI confidences interval
\*The red indicates observed values that fall within the 95% confidence interval, and the blue indicate values that are lower than predicated.

**Table 3.**
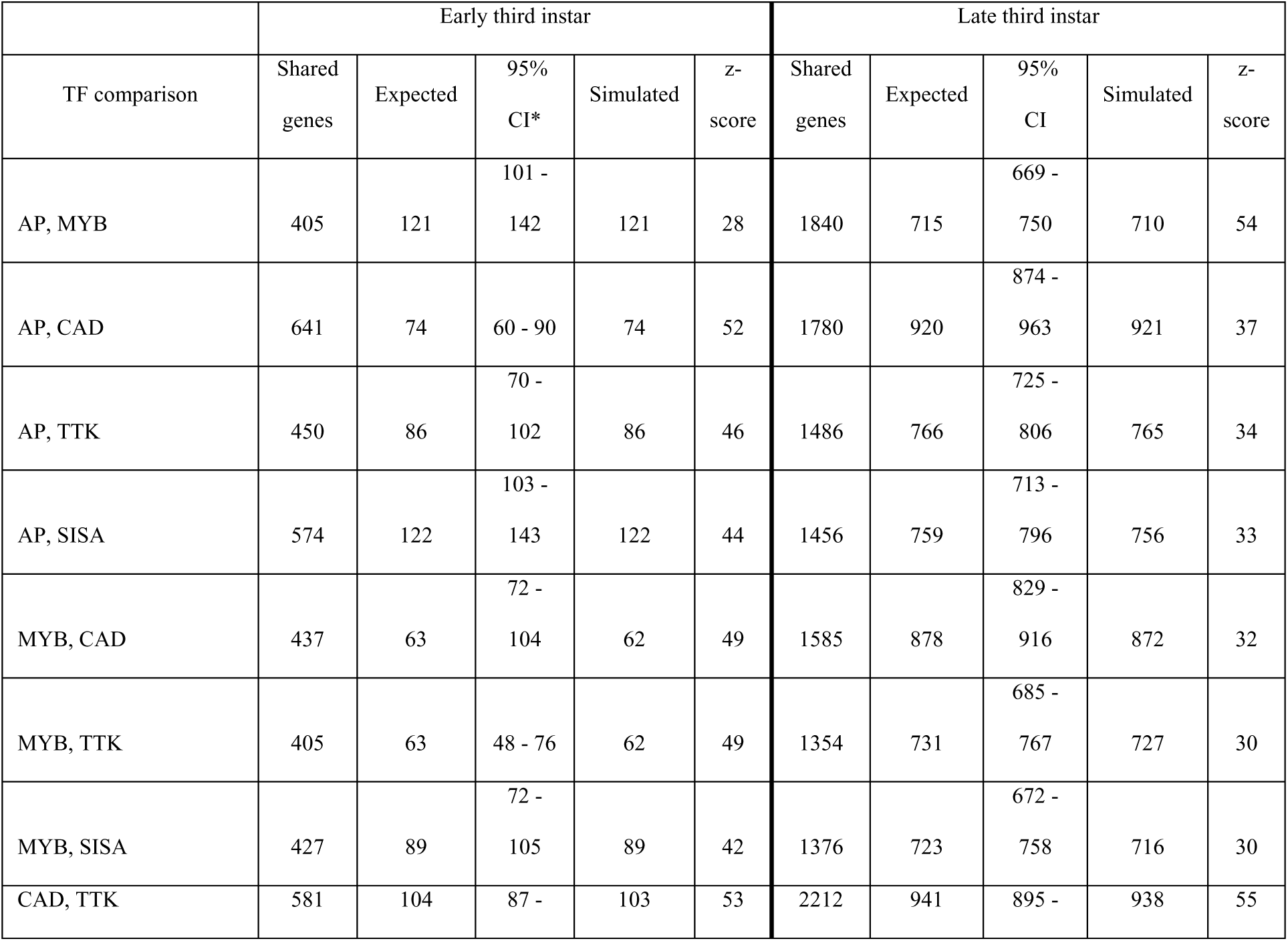
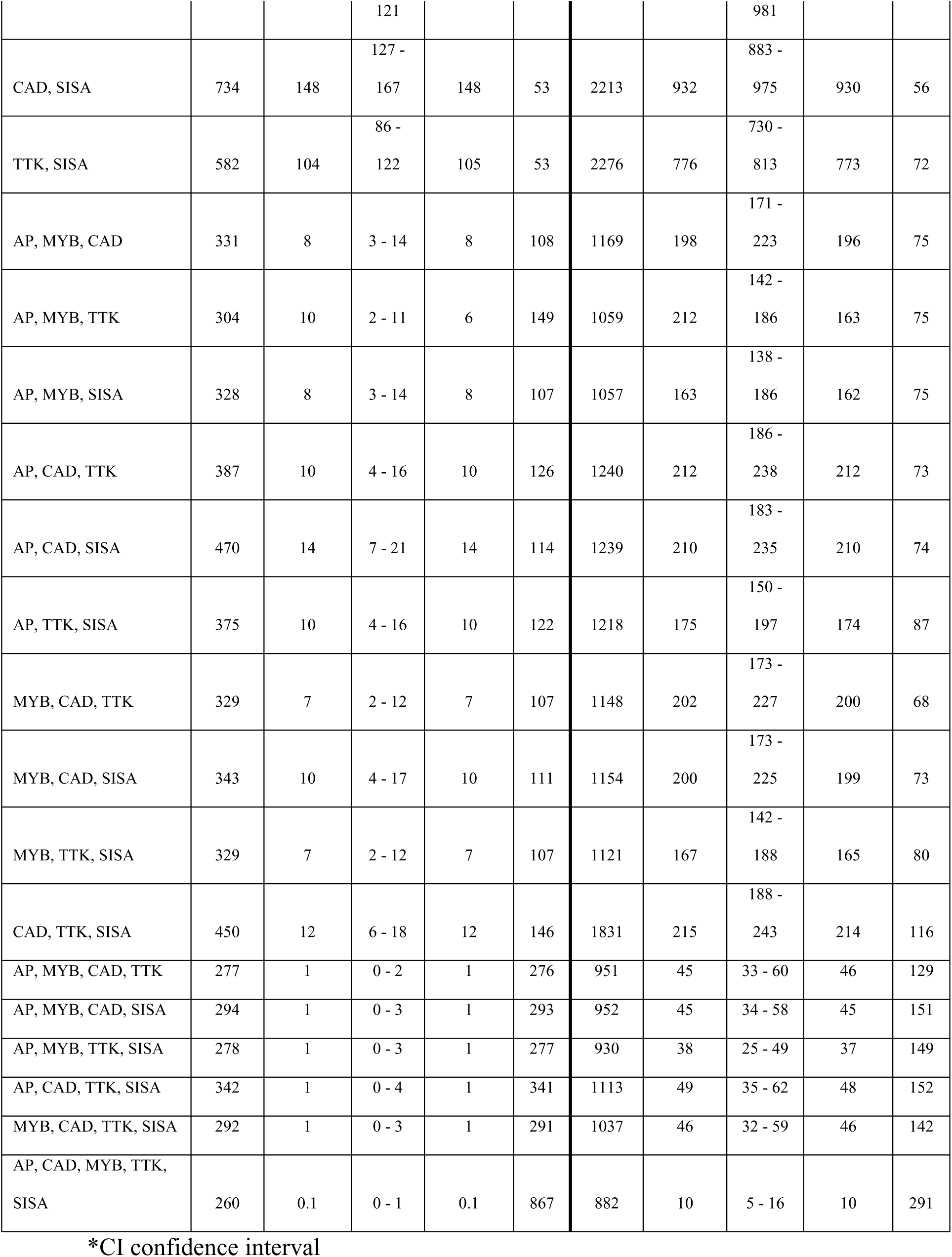
Non-independence of the overlap of transcriptomes in the rescue and non-rescue of ap.

Second, the overlap in a Venn diagram only represents a set of differential mRNA accumulation shared between two or more TFs and provides no information on the mode of regulation (positive or negative) and magnitude of the effect. Therefore, to characterize the transcriptomes further, the correlation between the effect of the TFs on log_2_ fold change (L2FC) of mRNA accumulation was assessed. Since in any analysis of differential mRNA accumulation about 50% are either up or down regulated by the action of a TF, there is no *a priori* reason to expect that two TFs have the same effect on the mode of regulation or the magnitude of regulation. The only other imposed factor on how mRNAs shared in a Venn diagram will behave in this analysis is that the shared mRNAs of both TFs (red/burgundy points) are required to have a strong enough effect on transcript accumulation to pass the *Padj* of <0.05 threshold for being a DEG. For stochastic behaviour, these considerations should result in little correlation between the L2FC of the two TFs, and the red/burgundy points, which are the shared mRNAs, should align in a diagonal cross with very few points along the x and y axes. This behavior was found for ANTP and BAB1 (Figure 2f; the only pair in Table-2 that falls within the simulated 95%CI); however, for most comparisons there was a positive correlation (Figure 2 g). When the Pearson’s correlation was calculated for all mRNAs dependent on either one TF and another TF (red/burgundy, green, magenta and blue points), the correlations were generally good with slopes approaching one indicating a similar magnitude of effect of both TFs (Figure 2e). The correlations were stronger when only the mRNAs that were dependent on both TFs were considered (red/burgundy points). Two interesting observations are: SQZ, which does not result in a female pattern of pigmentation, shows a strong correlation of transcript accumulation with BAB1 (Figure 2g), but ANTP, which does result in female-like pigmentation, shows little correlation (Figure 2f). Overall the correlations of transcript accumulation between TFs are not associated with phenotypic outcomes: FOXO and SQZ have the strongest correlations with BAB1, followed by ODD, then EY and finally ANTP but FOXO and SQZ do not rescue and ODD, EY and ANTP do. Both the simulation of stochastic behavior and analysis of correlations between transcriptomes suggest that the overlapping patterns of transcript accumulation are not stochastic and independent.

### Accumulation of mRNAs associated with sexually dimorphic pigmentation of the abdomen

The proteins encoded by the genes *y* and *t* are required for pigmentation of bristles and cuticle. Female cuticular pigmentation changes represent a small decrease in overall pigmentation of the abdomen. The accumulation of *Y* and *T* mRNAs overall (17/18) showed no changes (*P*>0.05) in the levels of accumulation relative to the *dsx^1^/dsxGAL4* control. The only exception was a decrease of *T* accumulation in males which is unexpected because male abdomens are more pigmented than female abdomens (Figure 3a). Although the requirement and regulation of *y* and *t* expression for cuticular pigmentation is well-characterized and established (Roeske et al., 2018), a potential role for Y and T in sexually dimorphic pigmentation would not have been identified in an RNAseq analysis of P13-15 male and female pupal abdomens, which is the time that Y and T are expressed and presumably required for abdominal pigmentation.

**Figure 3.**
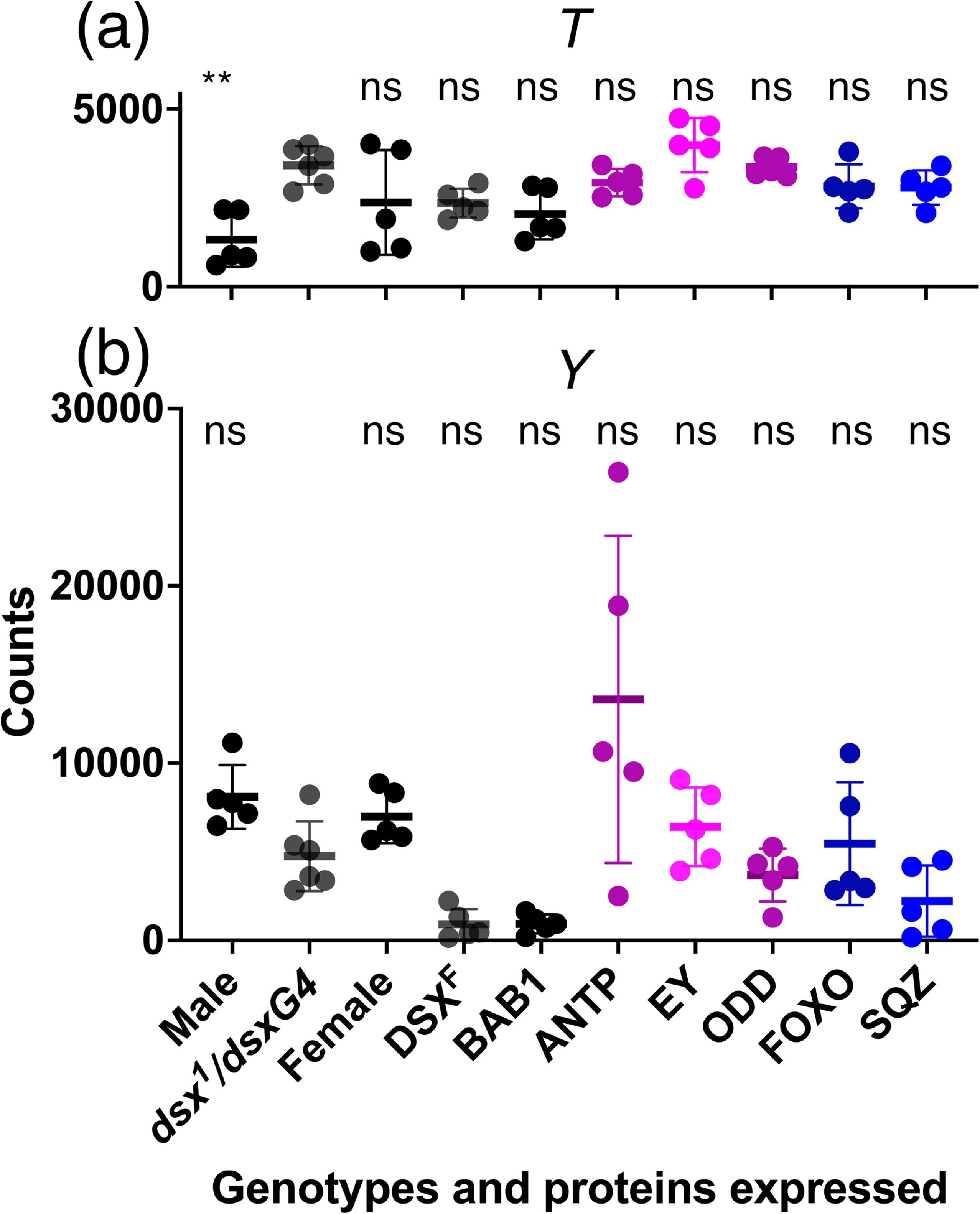
Accumulation of *T* and *Y* mRNA. Panels a and b are scatter plots of the normalized counts of *T* (a) and *Y* (b) mRNA accumulation for males, females, *dsx^1^/dsxGAL4* either not expressing a TF or expressing the TF indicated. At the top of the graphs is indicated the significance of any changes relative to *dsx^1^/dsxGAL4* in an ANOVA of ranks (*T* H(10) = 26, *P*<0.002; *Y* H(10) = 34, *P*<0.0001). The symbols: ns not significant; ** *P*<0.01.

### Temporal characterization of AP-dependent mRNA accumulation

During the second to early third instar larval stages, expression of AP initiates a pathway required for wing development (Figure 4a; Fisher et al., 2017). At these early stages of imaginal disc development, AP regulates the expression of FNG and SER which result in the activation of WG expression along the future wing margin (Diaz-Benjumea and Cohen 1993; Diaz-Benjumea and Cohen 1995; Irvine and Wieschaus, 1994). During the following 48 hours, WG patterns growth and development of the wing pouch. Larval wing imaginal discs are only large enough to dissect individually at the late third instar larval stage which is two days after the requirement of AP for initiation of wing development. To assess whether the AP-dependent transcriptome could be characterized closer to when AP is required, early third instar larvae were dissected by cutting them in half and removing the mouth hooks gut and brain to leave a patch of epidermis with leg, wing and haltere discs attached. AP is expressed and required in both the wing and haltere imaginal discs (Cohen et al., 1993). Comparison of the AP-dependent transcriptomes of early third instar larvae and late third instar larvae showed significant overlap in AP dependent mRNA accumulation in a Venn diagram (Figure b) and showed a good correlation when the log2 fold changes of AP-dependent mRNA accumulation at early and late third instar larval stages were compared (Figure 4c). Like the comparison of the DSX^F^ and BAB1 transcriptomes, the 484 genes that are shared in the Venn diagram is likely an underrepresentation as there are genes that were identified at one time point and show a similar log_2_ fold change in accumulation at the second time point but that did not meet the *P_adj_* <0.05 threshold at the second time point (blue and green points). However, the slope of the correlation is 0.6 (less than the expected 1) which is due mainly to a set of 13 points that have very high negative fold changes at E3 (-25 to -30) pulling the slope down (removing these points the slope is close to 1). These 13 genes all have high variability in the count data of the *ap^null^/apGAL4* control replicates which due to the methodology employed by DEseq2 to normalize the data and shrink log_2_ fold changes inflates the fold change (Love et al., 2014). Correlation between AP-dependent transcriptomes at early third and late third instar stages suggests that similar sets of AP-dependent transcripts are identified at these distinct temporal stages even though the early third instar larval tissues are not composed of only wing disc cells and that AP is also expressed in the haltere imaginal disc. Therefore, analysis of TF-dependent transcriptomes associated with phenotypic nonspecificity at early third instar larval stage, a time close to AP requirement, is feasible.

**Figure 4.**
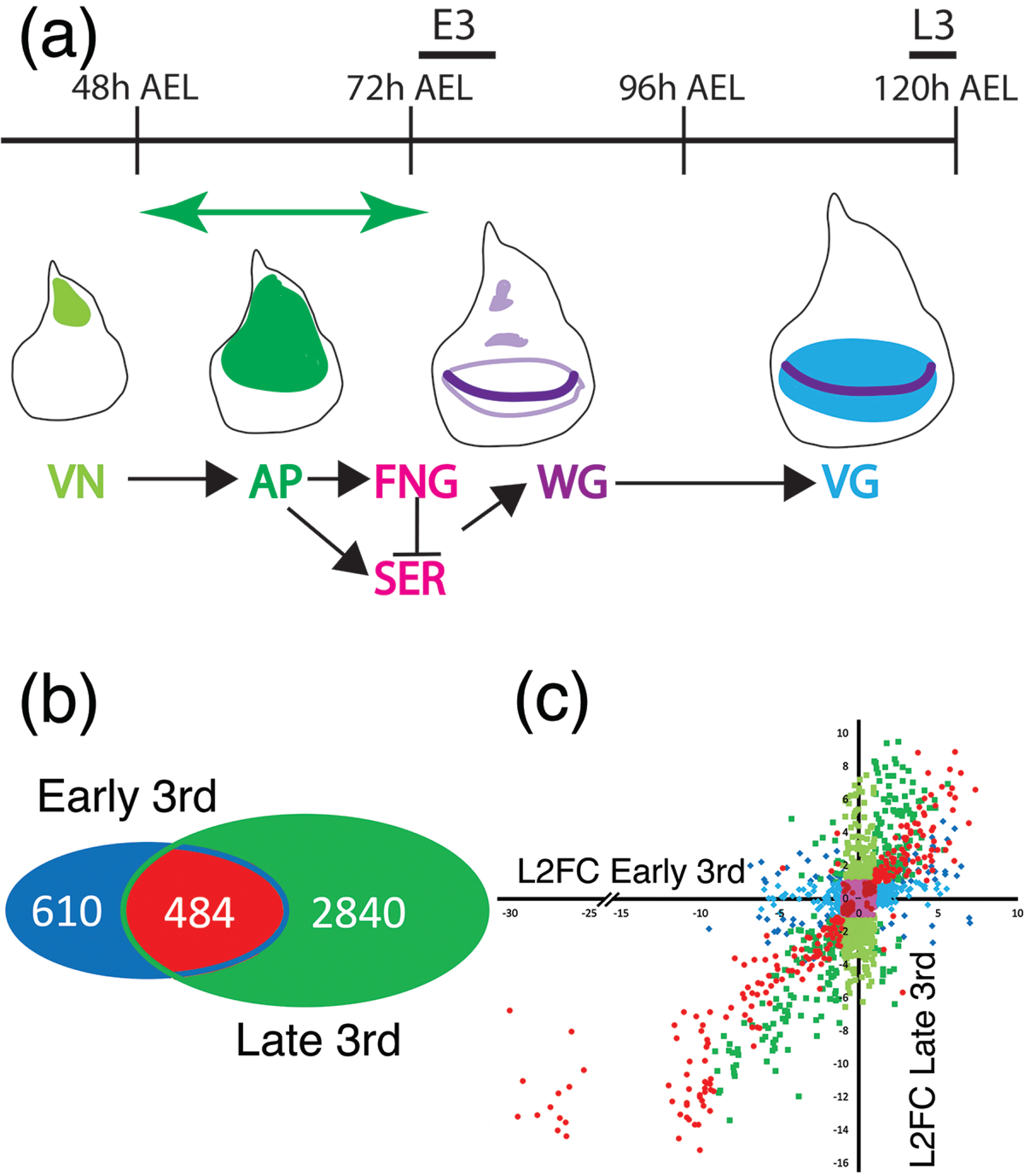
Temporal analysis of AP dependent mRNA accumulation during larval development. a. Representation of the time course of wing imaginal disc development during the larval stages (AEL after egg laying). Below the time are representations of developing and growing imaginal disc (the increase in size is not drawn to scale). During the first and early second instar larval stage VN and WG determine the dorsal ventral axis, and VN activates the expression of AP (green). During the second instar and into the early third instar larval stage (E3)(the green double-headed arrow), AP is required for wing formation by activating FNG and SER expression (Diaz-Benjumea and Cohen 1993; Diaz-Benjumea and Cohen 1995; Irvine and Wieschaus, 1994; Fisher et al., 2017). FNG and SER through the regulation of Notch activity establish the expression of WG in the cells of the future wing margin (dark purple) by the early third instar larval stage. During the next 40-48 hours till the late third instar larval stage (L3), the WG morphogen in cooperation with the DPP morphogen stimulate growth and differentiation of the wing pouch by regulating patterns of gene expression such as the expression of VG (blue). The bars above the time-line indicate the time intervals that tissue was dissected for RNA extraction. b. Venn diagrams of the AP-dependent transcriptomes identified by DEseq2 analysis at E3 and L3. c. Scatter plot of the log_2_ fold change (L2FC) of AP-dependent mRNA accumulation at E3 plotted on the x-axis versus AP-dependent mRNA accumulation at L3 plotted on the y-axis. The circular points in red and burgundy are DEGs identified in both the E3 and L3 transcriptomes as AP dependent (*P_adj_* < 0.05). The red points have a L2FC >1 and <-1; the burgundy points have a L2FC < 1 and > -1. The diamond blue/light blue points are DEGs identified in the E3 transcriptome (*P_adj_* <0.05) but not the L3 transcriptome (*P_adj_* >0.05). The blue points have a L2FC >1 and < -1 at L3; light blue have a L2FC <1 and > -1 at L3. The square green points are DEGs identified in the L3 transcriptome (*P_adj_* < 0.05) and not the E3 transcriptome (*P_adj_* > 0.05). The green points have a L2FC >1 and <-1 at E3; light green L2FC < 1; >-1 at E3. Magenta points were identified as DEGs in either the E2 or L3 transcriptomes with a L2FC of <1 and >-1 in both.

### TF-dependent transcriptomes associated with rescue/non-rescue of wing development

The ap wingless phenotype is partially rescued by expression of CAD, MYB or TTK, but not SISA (Percival-Smith et al., 2023). To characterize the rescue further, wing imaginal discs from *ap* mutant third instar larvae expressing AP, CAD, MYB, TTK or SISA were dissected and *WG* expression assessed. The wing discs expressing GAL4 alone or with SISA have a circular ring of WG expression in the reduced wing pouch (Figure 5a). In the wing discs expressing CAD, TTK or MYB, *WG* is expressed in a crescent shape that is potentially along the dorsal ventral boundary of the disc (Figure 5a). The expression of AP rescues the expression of *WG* in wing margin cells.

**Figure 5.**
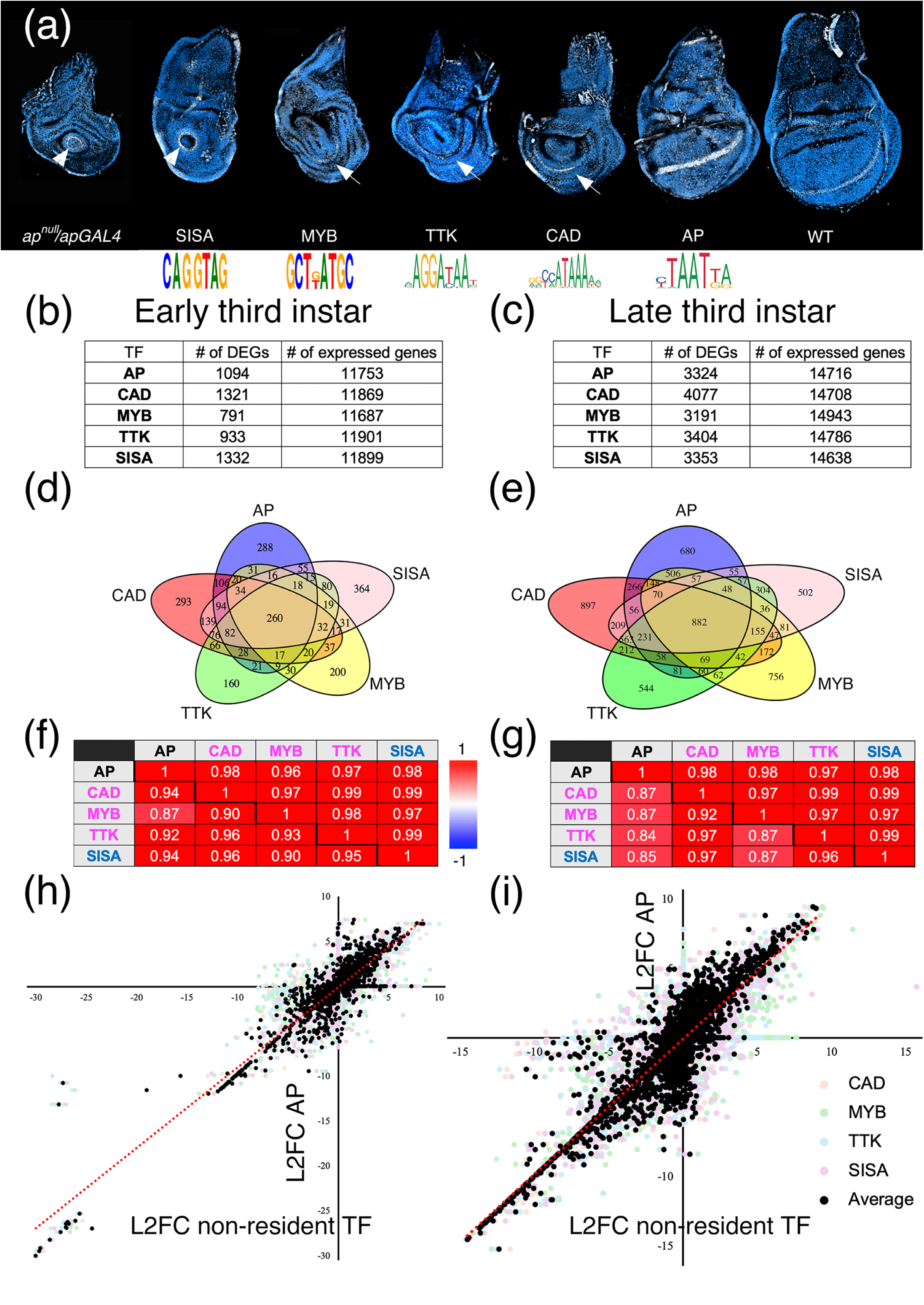
Analysis of transcriptomes associated and not associated with rescue of the wingless ap phenotype. a. *WG* in situ hybridization (white) on third instar imaginal discs with nuclei identified with DAPI (Blue). The genotype or protein expressed are indicated below. The arrowheads point to a circle of *WG* expression and the arrows point to a crescent of *WG* expression potentially along the dorsal ventral boundary. Below the discs expressing particular TFs is the sequence logo of the AP, CAD, MYB, and TTK TFs, and the consensus binding sequence for the SISA TF (Erikson and Cline, 1998; Vlieghe et al., 2006). b. Table of the number of DEGs and expressed genes (base mean>1) identified in the DEseq2 analyses at the early third instar larval stage. c. Table of the number of DEGs and expressed genes (base mean>1) identified in the DEseq2 analyses at the late third instar larval stage. d. Venn diagram of DEGs identified in *ap^null^/apGAL4* mutant background expressing AP, CAD, MYB, TTK and SISA at the early third instar larval stage. e. Venn diagram of DEGs identified in *ap^null^/apGAL4* mutant background expressing AP, CAD, MYB, TTK and SISA at the late third instar larval stage. f. Table of Pearson’s correlations between DEGs of various TF transcriptomes at early third instar larval stage. The diagonal separates two sets of analyses. The Pearson’s correlations below and left of the diagonal are for all the DEGs identified in either of the transcriptomes of the two TFs. The Pearson’s correlations above and right of the diagonal are for the DEGs identified in both transcriptomes (*P_adj_* < 0.05 for both). g. The same as f with the exception that they are the transcriptomes at the late third instar larval stage. h. Scatter plot of the log_2_ fold change (L2FC) of AP-dependent mRNA accumulation plotted on the y-axis versus the L2FC for CAD, MYB, TTK and SISA individually (lightly colored points, see legend), and the average of CAD, MYB, TTK and SISA (black points) dependent mRNA accumulation plotted on the x-axis for the transcriptomes at early third instar larval stage. i Same as g except the transcriptomes are at late third instar larval stage. The dotted red line in f and g is the tread-line. The legend for g and h is shown at the bottom right.

The TF-dependent transcriptomes were identified at both the early and late third instar larval stages. The Venn diagram comparison of the TF-dependent transcriptomes (AP, CAD, MYB, TTK and SISA) at both larval stages showed extensive overlap of TF-dependent mRNA accumulation: at early third instar larval stage accumulation of 1,373 mRNAs were dependent on two or more TFs and 1,305 were dependent on a single TF; at late third instar larval stage accumulation of 3,467 mRNAs were dependent on two or more TFs and 3,379 were dependent on a single TF. Using the number of DEGs and the total number of expressed mRNAs to simulate the expected values and determine a 95%CI based on stochastic behaviour, not a single observed value was low enough to fall into the CI and all z-scores were high. When all the L2FCs of the mRNAs that changed in one of two TF analyses were plotted on either the x or y axis, the correlations where high. The correlation increased when only those shared between the two TFs were compared (Figure 5d,e). The correlations were so high at both stages that the average of the fold changes observed with CAD, MYB, TTK and SISA for 2,678 mRNAs at E3 and 6,846 mRNAs at L3 were plotted against the fold change observed with AP (Figure 5f, g; black points). At the early third instar stage 90% of the four averaged TFs had all four values either all positive or all negative and this increased to 96% at the late third instar larval stage. The slope of the correlation was close to 1 indicating that the magnitude of the effect on mRNA accumulation was similar with all TFs. To indicate the variation observed, the fold changes of the individual TFs CAD, MYB, TTK and SISA were also plotted against those of AP (Figure 5 f,g lightly colored points). One striking set of points are the near perfect line of points in the bottom left quadrant of the graphs comprised of genes repressed by all TFs. These points all have counts in the control replicates and very low or zero counts with any of the TFs expressed. This pattern of counts in the data, in combination with the similarity between overall transcriptomes, results in the very similar L2FC with different TFs producing a straight line of points. The transcriptomes associated with the resident AP TF and these four non-resident TFs show strong co-expression suggesting the regulation of a constrained set of genes. The correlation was strong irrespective of whether the ap phenotype was rescued or not.

### Genes potentially directly regulated by AP

At the late third instar larval stage the total number of mRNAs showing a highly corelated change in accumulation is very high at close to 40% of the expressed mRNAs. Therefore, it is possible that AP and the non-resident TFs (CAD, MYB, TTK and SISA) trip a common switch that initiates a massive common response that obscures the specific and diverse regulation of genes by these five different TFs within the dorsal AP-expressing cells (Figure 6a). Our approach to address this issue was to ask whether mRNAs expressed in the dorsal epithelium of the wing imaginal disc had been identified as AP-dependent. The virtual in situ tool developed by Everett et al., 2021, was used to classify all the AP-dependent mRNAs into 11 classes of expression patterns (Figure 6b-l). No obvious preference was observed in how the 951 mRNAs shared by AP, CAD, MYB and TTK partitioned between the 11 expression categories; each category have about 25% of these mRNAs. The 146 AP-dependent mRNAs expressed in dorsal cells and 147 that were identified as non-resident TF-dependent and expressed in dorsal cells were plotted AP versus the average of the four non-resident TFs (Figure 6m,n; black circles) and AP versus the individual TFs CAD, MYB, TTK and SISA (Figure 6m,n the lightly colored points). The correlation between these mRNA expressed in the AP-expressing cells, and therefore, potentially directly regulated by AP was very high at both the late and early third instar larval stage. Although not all of these 294 (4% of the total DEGs) genes may be directly regulated by AP, there was still a strong correlation of expression observed allowing the average of the four non-resident TFs to be plotted against AP, which would not have been the case if diverse gene regulation patterns were initiated in dorsal cells by the different TFs.

**Figure 6.**
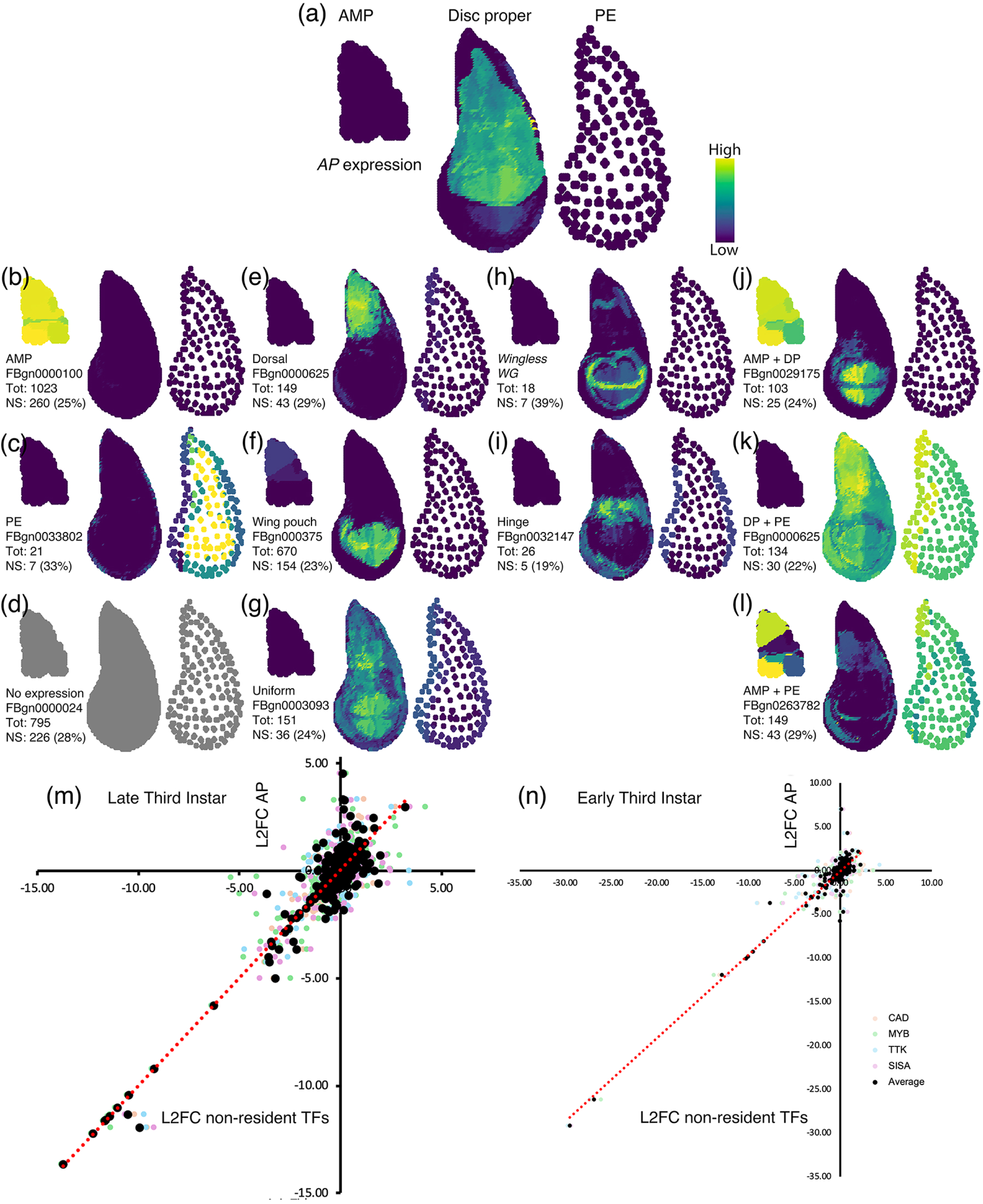
Virtual in situ analysis of AP-dependent mRNA accumulation and the change of AP-dependent mRNA accumulation in AP-expressing cells. a A virtual in situ of AP expression in the dorsal compartment of the disc proper. The expression in all in situs are shown in the mesodermal adult muscle precursor cells (AMP), the epithelial disc proper cells and the peripodial envelope cells (PE). b-l In situs for all 3,324 AP dependent mRNAs sorted into 11 expression categories of which an exemplar is shown. The total (Tot) number of mRNAs in each non-overlapping category is given. The number of mRNAs shared by AP, CAD, MYB and TTK are given below the total (Phenotypically nonspecific: NS) for each category with a percent of the total given. m. Scatter plot of the log_2_ fold change (L2FC) of AP-dependent mRNA accumulation plotted on the x-axis versus the average of CAD, MYB, TTK and SISA dependent mRNA accumulation plotted on the y-axis for the transcriptomes at late third instar larval stage for the 294 genes that are expressed in the dorsal compartment (black circles). Along with the average are plotted the AP vs the four individual TFs (CAD light orange, MYB light green, TTK light blue and SISA light red) to display the variance in the data. n. Scatter plot is as shown in m but with data from the early third instar larval stage (base mean >1).

### Expression of genes required for wing development

The WG developmental pathway required for wing development is initiated by the AP TF. AP regulates the expression of two genes, *frn* and *Ser*, whose products regulate Notch receptor activity required for establishment of WG secretion from the cells of the future wing margin. The WG morphogen activates the expression of the *vg* gene. The components of this pathway were identified by the phenotype of mutations; therefore, would RNAseq analysis at late third instar larval stage have identified this developmental pathway? The *FNG* mRNA is present in all samples but is unexpectedly lower in wild type imaginal wing discs (Figure 7a). The *SER* mRNA does increase in abundance in WT and AP rescued wing discs (Figure 7b). The *WG* mRNA does increase in abundance in WT, and AP rescued wing imaginal discs as defined by having a *P*<0.05 (Figure 7c), but it should also be noted that the magnitude of *WG* expression was higher in WT, AP, CAD, TTK and MYB than in both *ap^null^/apGAL4* and SISA. However, these 3 mRNAs do not meet the threshold of a 2-fold change set in most RNAseq analyses for the differential expression effect size and would have been overlooked. This small change is expected from prior expression analyses: at the late third instar larval stage *FRN* mRNA is uniformly expressed across the wing imaginal disc; expression of SER protein is increased in the AP domain but not restricted to the AP domain of early third instar larval imaginal discs; in wing imaginal discs WG expression is not restricted only to the cells of the future wing margin (Irvine and Weischaus, 1994; Diaz-Benjumea and Cohen, 1995; Calvo et al., 2021). *VG* mRNA strongly accumulates in WT and AP rescued wing imaginal discs, but not in CAD, MYB, or TTK rescued wing imaginal discs (Figure 7d). Therefore, only *VG* would have been identified in an RNAseq analysis.

**Figure 7.**
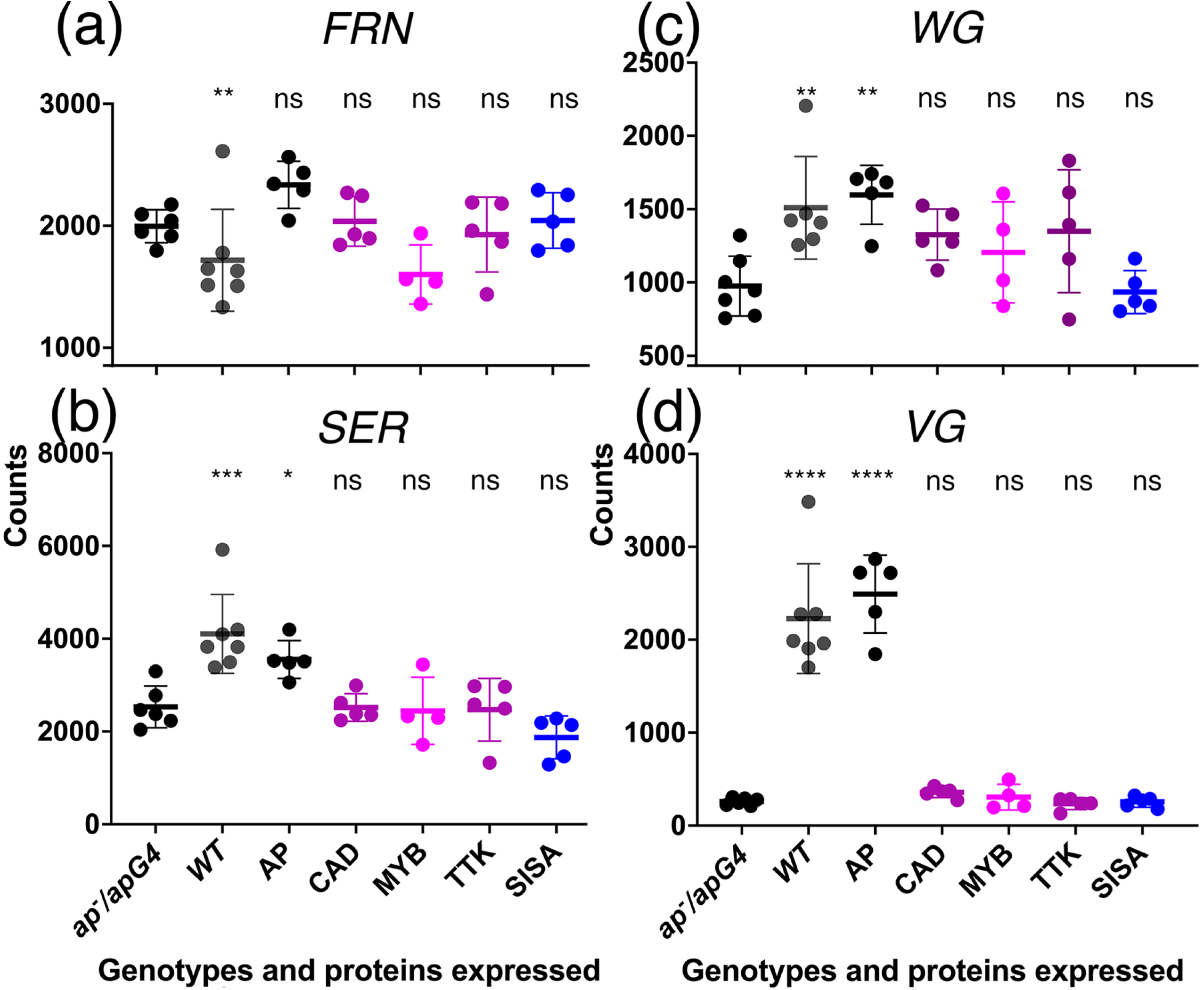
Accumulation of *FRN, SER, WG* and *VG* mRNA. Panels A-D are scatter plots of the normalized counts of *FRN* (A), *SER* (B), *WG* (C) and *VG* (D) mRNA accumulation in *ap^null^/apGAL4* either not expressing a TF or expressing the TF indicated or WT. At the top of the graph is indicated the significance of any changes relative to *ap^null^/apGAL4* in an ordinary ANOVA (*FRN* F_7, 33_ = 6, *P*<0.0002; *SER* F_7, 33_ = 9, *P*<0.0001; *WG* F_7, 33_ = 5, *P*<0.001; *VG* F_7, 33_ = 64, *P*<0.0001). The symbols: ns not significant; * *P*<0.05; ** *P*<0.01; *** *P*< 0.001; **** *P*<0.0001.

## Discussion

### Transcriptomes associated with phenotypic nonspecificity

The goal that initiated this study was distinguishing between three possible expectations for the transcriptomes associated with phenotypic nonspecificity. The transcriptomes associated with TFs that both rescued and did not rescue the dsx and ap phenotypes were overlapping sets of mRNA accumulation that were not stochastic/independent. The lack of independence was observed as a higher than expected overlap between transcriptomes and a high correlation of the responses to expression of different TFs. For both the dsx and ap systems, the regulation of a constrained set of genes is supported. However, these correlations in response to expression of different TFs are not good predictors of phenotypic nonspecificity. The ANTP transcriptome, which shows low correlation with BAB1 transcriptome, does rescue but the SQZ transcriptome, which corelates well with the BAB1 transcriptome, does not rescue. For the rescue of the ap phenotype, the SISA transcriptome, which does not rescue the ap phenotype, does have a high correlation with the AP transcriptome. The high degree of correlation between the TF-dependent transcriptomes would have led to speculation that some of the non-resident TFs may be able to rescue the dsx or ap phenotype, if this transcriptomic study had been performed without knowledge of phenotypic nonspecificity.

### Proposed mechanisms for phenotypic nonspecificity

The systematic analyses of TF function and real-time visualization of transcription has challenged many common assumptions of the role of eukaryotic TFs in regulating the rate of transcription initiation. These four assumptions are: i) a primary requirement for the DBD in occupying particular sites in the genome; ii) clear modularity/separation of domains required for occupancy, transcriptional regulation and pleiotropy; iii) mutual cooperativity in occupancy; and iv) assembly of TFs and RNA polymerase results in a constant rate of transcriptional initiation. A large-scale analysis of the requirement of the DNA-binding domain (DBD) of 48 yeast TFs for genome occupancy concluded that the mechanism of occupancy of TFs can be anywhere on a spectrum from complete DBD-dependence to complete DBD-independence (Brodsky et al., 2020; Kumar et al., 2023). The DBD-dependence/independence of the Drosophila Fushi tarazu (FTZ) TF is differentially pleiotropic with FTZ being DBD-independent for embryonic segmentation but DBD-dependent for deletion of wings and eyes suggesting that where TFs sit on the DBD dependence/independence spectrum can change both temporally and spatially (Hyduk and Percival-Smith 1996; Percival-Smith, 2017). Systematic mutational analysis of the yeast Msn2p TF has uncovered two characteristics of occupancy and transcriptional activity. First, occupancy does not require the DBD and instead is dependent on intrinsically disordered regions (IDR) of the protein. Second, there are amino acids important for transcriptional activity but not for occupancy in the IDR, indicating that binding and expression determinants in the IDR are dispersed, separable and intermingled. These determinants important for both binding and expression are short dispersed elements and sequences of low complexity (Mindel et al., 2024). The analysis of Msn2p was performed in a single cell type and with the same environmental conditions but analysis of TF function/pleiotropy in different tissues/phenotypes indicates that the dispersed functional elements of TFs outside the DBD make small differential contributions to overall TF activity (Bhoite et al., 2002; Hittinger et al., 2005; Sivanantharajah and Percival-Smith, 2009, 2014, 2015; Percival-Smith et al., 2013), as opposed to TFs being composed of compact domains with specific requirements for TF activity in a particular tissue. In addition, differential pleiotropy of TF function extends to the coordinate regulation of gene expression within a cell. In a yeast cell, Bas1p and Bas2p are two TFs required for expression of *HIS4* and *ADE5,7* genes, and the requirement of Bas2p amino acids for expression of *HIS4* and *ADE5,7* differ providing an example of differential pleiotropy within the same cell (Bhoite et al., 2002). Cooperativity in the co-occupancy of a set of 12 TFs associated with the yeast stress response is not mutual but one sided: Msn2p binding is required for the binding of the other 11 TFs; however, the other 11 TFs are not required for the binding of Msn2p to its sites (Lupo et al., 2023). The initiation rate of an actively transcribed gene observed in real time is pulsed and not continuous (Chubb et al., 2006). Analysis of the TF clusters that form, and believed important for transcription bursting, are dynamic assemblies (Rodriguez and Larson, 2020). These observations suggest that the traditional view of how TFs act to regulate the rate of transcription requires revision at the very least.

The limited specificity of TF function and the assembly of TFs into membrane-less compartments, wolfpacks, are distinct models used to explain phenotypic nonspecificity (Percival-Smith, 2018; Staller, 2022; Percival-Smith et al., 2023). Limited specificity proposes that TFs are independent entities that bind many sites in the genome using relatively non-specific cooperative interactions and transcription is an emergent property associated with the number of TFs bound to an enhancer/silencer. In this model TFs regulate distinct patterns of gene expression, and phenotypic nonspecificity is a consequence of the overlap between the distinct patterns of gene expression containing the information required for rescue of the phenotype. In this unbiased model for phenotypic nonspecificity, the transcriptomes of TF-dependent mRNA accumulation should behave stochastically relative to one another which is not supported by the analyses of dsx and ap sets of transcriptomes. The wolfpack along with the lone wolf model for TF genome occupancy explain DBD independence observed with many yeast TFs (Brodsky et al., 2020; Staller, 2022; Kumar et al., 2023). To explain phenotypic nonspecificity, the wolfpack model was modified such that each nucleus contains a discrete number of wolfpacks containing a similar number of TFs that is proportionally related to the frequency of phenotypic nonspecificity. For Drosophila the estimated number of wolfpacks in the nucleus is 10-20 based on a frequency of phenotypic non-specificity of 1 in 10-20 TFs, and therefore, if every cell expresses between 100-150 TFs each wolfpack is composed of between 5-15 TFs (Percival-Smith et al., 2023). As each TF in a wolfpack makes a small contribution to the information required to find specific binding sites in a genome, the substitution of one TF (TFa) for another (TFb) has little consequence on the sites bound in the genome because the information provided by the other 4-14 TFs dominates over that of TFa and TFb for the sites bound such that wolfpacks containing either TFa or TFb share a high percentage of sites (Wunderlich and Mirny, 2009). The transcriptomes associated with the expression of TFa and TFb should be similar due to the regulation of a constrained set of genes as observed with most transcriptomes in this study. The clear exception to this is the lack of correlation in the dsx set between the ANTP transcriptome and the other transcriptomes. This lack of correlation is potentially explained by ANTP participating in two or more distinct wolfpacks obscuring correlation which would also be consistent with the observation that ANTP has the largest number of DEGs relative to the other TFs studied in the dsx set.

The nucleoplasm is separated into many specialized nonmembrane bound compartments: the nucleolus required for rRNA transcription and transcript processing; histone locus bodies for replication dependent expression of histone mRNAs; and Cajal bodies (Liu et al., 2006; Falahati et al., 2016; Geisler et al., 2023). The wolfpack model, as used to explain phenotypic nonspecificity, suggests that the 100-150 TFs expressed in a Drosophila cell are partitioned into 10-20 nucleoplasm compartments, and when a compartment interacts with a gene’s regulatory DNA sequence may initiate the formation of a transcription hub. An expectation of this proposal is the genome occupancy of TFs from the same wolfpack would correlate. Analysis of 141/150 yeast TFs expressed during partial nutrient deprivation show examples of colocalization/correlation: 23 TF associated with Cyc8; 12 TF associated with Med-15; plus 14-18 groups composed of 2-9 TFs; but 50-60 TFs do not corelate strongly with others (Lupo et al., 2023). Although, about two thirds of yeast TFs show correlation in binding, yeast do not contain 10-20 neat and tidy TF compartments of approximately equal size; however, due to the small size of yeast regulatory sequences relative to those of other eukaryotes, it is suggested that the frequency of phenotypic nonspecificity would be lower in yeast in turn suggesting a larger number of nuclear compartments exist in yeast (Percival-Smith, 2018).

Although the wolfpack model explains the constrained set of genes regulated by TFs observed, another potential explanation is that TFs bind to regulatory sites after chromatin has been made accessible. Since eukaryotic TFs recognize sequences of low information content, any region of the genome opened up and accessible would be bound by both TFa and TFb explaining the DNA-binding site recognition independence of phenotypic nonspecificity (Wunderlich and Mirny, 2009). One model proposed for creating accessible chromatin is that a specialized class of TFs, pioneer TFs, bind to DNA in nucleosomes and open up chromatin allowing access to DNA by other non-pioneer TFs (Zaret, 2020). This model has some experimental support and also an exception (Hansen et al., 2022; Brennan et al., 2023). A remaining problem is how pioneer TFs, which themselves recognize DNA binding sites with low information content, find specific sites in the genome to initiate making chromatin accessible (Wunderlich and Mirny, 2009). It is possible that in formation of a wolfpack, pioneering function is imparted to the wolfpack through incorporating a subset of TFs that can interact with DNA in nucleosomes providing entry of the whole wolfpack to specific sites in chromatin.

### Transcriptomes and phenotypes

A common assumption made about the regulation of gene expression and its association with phenotype is that trans-acting regulatory factors active in a cell activate the expression of a specific set of RNAs (transcriptome) and proteins (translatome) that then fashion the observed phenotype. This assumption is supported at the individual gene level where removal or addition of a function expressed in a restricted temporal and spatial pattern does change phenotype; however, the assumption is only now being tested experimentally at the level of the genome, transcriptome and translatome. Both phenotypic nonspecificity and phenotypic convergence suggest that unique sets of TFs bring about similar transcriptomes or phenotypes (Konstantinides et al., 2018; Percival-Smith et al., 2023). “You show me which genes are on, I’ll tell you what cell type” is a central tenet important for the interpretation of both scRNAseq and RNAseq analyses (Hahn et al., 2023; Dance, 2024). Indeed, in the *C. elegans* nervous system, the transcriptome type (t-type) of neurons share morphology (m-type) and function (f-type); however, in the zebrafish optic tectum, single t-types can have more than one m-type and f-type (Taylor et al., 2021; Shainer et al., 2025). We found that there is not a strong correlation between the overall structure of the transcriptomes and the phenotype generated. This is similar to the intraspecies differences in developmental transcriptomes expressed during sporulation of yeast. Two yeast strains share only 60% of the genes differentially expressed during sporulation (Primig et al., 2000) but both nonetheless sporulate. Although these observations require more examples in a diverse range of species, it should be taken into account in interpretation of any future analyses, as transcriptomes are being used as markers of cancer cell type, which may require more investigation for use as a precise diagnostic tool (Vareslija et al., 2019; Tsimberidou et al., 2022).

RNAseq is inefficient at identifying genes important for a process. In our RNAseq analyses, many genes known to be involved in abdominal pigmentation and wing formation would not have been identified. The systematic functional genomic analysis of yeast sporulation provides two other issues with RNAseq analyses: of the many genes expressed preferentially during sporulation only a low proportion are required, and of the genes identified by a non-lethal deletion required for sporulation, only 39% are upregulated during sporulation and most surprisingly 21% of the genes required are down regulated (Deutschbauer et al., 2002; Enyenihi and Saunders, 2003). Most RNAseq analyses screening for candidates that are required for a particular process ignore downregulated genes. Although some, or much, of the inefficiency observed in yeast sporulation may be due to extensive redundancy, as has been observed during vegetative growth (Costanzo et al., 2010), the poor intraspecies conservation of the transcriptional program associated with sporulation points to major deficiencies in RNAseq analyses for efficiently identifying factors important for developmental processes. A major deficiency in using RNAseq to identify genes regulated directly by the binding of TFs to the gene’s regulatory sequence in a multicellular organism composed of many cell types is highlighted in this study by only 4% of AP-dependent mRNA accumulation being assigned to AP expressing cells.

### Limitations and future studies

Although this study accomplishes its initial goal of distinguishing between three general transcriptomic outcomes, it can only speculate on the underlying mechanisms that govern the expression of similar sets of genes by multiple TFs. From the point of view of the study of phenotypic nonspecificity, the dsx and ap phenotypic systems are good in that multiple examples of nonspecificity were identified, but from the perspective of transcriptome analysis, the abdomen and wing imaginal disc are complex organs composed of many cell type specific transcriptomes preventing more far reaching conclusions. For future studies of the mechanism that explains phenotypic nonspecificity, the system employed must have multiple examples of phenotypic nonspecificity. But to thoroughly investigate whether a limited number of wolfpacks and expression programs exist in cells expressing the TF, these cells must have one cellular/transcriptomic phenotype and can be isolated in large enough numbers during determination to perform an analysis of occupancy of all the TFs expressed in the cells.

## Data Availability

The raw clean forward read data and extracted count data used for analyses is available on the Gene Expression Omnibus (GEO) at GSE296804.

## Acknowledgements

We thank Ben Rubin for assistance with the statistical analysis and the Monte Carlo simulation in particular. We thank Greg Gloor for advice on our RNAseq analyses. We thank Paras Gupta for assistance in mRNA extraction for some of the samples. The extended focus imaging of the abdomens and confocal microscopy were performed at the Biotron Integrated Microscopy Laboratory (London, ON), and thank Karen Nygard and Amanda Casselman for assistance in the imaging of the samples.

Contributions to the paper: AP-S designed the study. AP-S constructed the genetic stocks. AP-S and GS performed the dissections and extracted the mRNA. AP-S performed the *WG* in situ hybridization. GS analyzed all the transcriptomics data and uploaded the data to the GEO site. AP-S wrote the first draft of the manuscript and AP-S and GS revised the drafts.

## Alt Text for Figures

Figure 1. The Venn diagram and scatter plot of log2 fold change of DSX^F^ (x-axis) and BAB1 (y-axis) regulated genes showing that BAB1 regulated genes are a sub-set of DSX^F^ regulated genes.

Figure 2. The Venn diagrams and log2 fold change plots of DSXF, BAB1, ANTP, EY, ODD, FOXO and SQZ regulated genes. A Are the abdominal phenotypes of the rescue of female-like pigmentation by DSXF, BAB1, ANTP, EY, ODD but not FOXO or SQZ. B. Table of the number of regulated and total expressed genes. C and D. Venn diagrams of BAB1, ANTP, EY and ODD regulated genes and BAB1, ANTP, EY, FOXO and SQZ regulated genes showing overlapping sets of genes. E. Correlations observed between TF regulated transcriptomes. F and G. Two log2 fold change scatter plots of BAB1 regulated genes (y-axis) and either the ANTP regulated genes or SQZ regulated genes (x-axis).

Figure 3. Scatter plots of the counts of the *Tan* and *Yellow* transcripts in the abdomens of males, *dsx^null^* mutants, females, and *dsx^null^* mutants expressing DSXF, BAB1, ANTP, EY, ODD, FOXO or SQZ showing no significant changes in expression with one unexpected exception.

Figure 4. Identification of AP regulated genes at the early and late third instar larval stage. A is a diagram of the timeline of the pathway required for wing development initiated by AP expression in the dorsal compartment. B Venn diagram of the overlap of early and late third instar AP regulated genes. C is scatter plot of log2 fold change of early (x-axis) and late (y-axis) third instar AP-regulated genes showing correlation.

Figure 5. The Venn diagrams and log_2_ fold change plots of AP, CAD, MYB, TTK and SISA regulated genes showing extensive overlapping expression. A show the rescue of *WG* expression patterns in *ap^null^*mutants not expressing a TF or expressing either SISA, CAD, MYB, or TTK. B, C are the table of DEGs and total expressed genes at the early or late stage. D, F are the Venn diagrams of the TF regulated genes at the early or late stage. G, H are the correlations between the TF-regulated genes at the early and late stage. I and J are the log_2_ fold change scatter plots of AP-regulated genes (x-axis and the average of CAD, MYB, TTK and SISA regulated genes (y-axis) at the early and late stage.

Figure 6. The virtual in situs of AP-regulated genes and scatter plots of log2 fold changes of genes expressed in the dorsal wing disc proper. A shows a virtual in situs for AP-expression followed B-L by examples of an expression pattern representative of a type of expression. M and N are the log_2_ fold change scatter plots of AP-regulated genes (x-axis and the average of CAD, MYB, TTK and SISA regulated genes (y-axis) at the late and early stage.

Figure 7. Scatter plots of the counts of the *FNG, SER, WG* and *VG* transcripts in the wing discs of *ap^null^* mutants, wild-type, and *ap^null^* mutants expressing AP, CAD, MYB, TTK or SISA showing that only *VG* would have been identified in a RNAseq analysis.

